# Mapping the distribution of elms in Britain after a century of Dutch elm disease

**DOI:** 10.64898/2026.07.16.738901

**Authors:** Catherine Walter, Mohammad Vatanparast, Camila Quintero-Berns, David Shreeve, Clive Brasier, Joan F. Webber, Richard J. A. Buggs

## Abstract

The establishment of novel pests and pathogens that devastate populations of trees is increasing around the world, giving rise to policy and management decisions about how to restore treescapes. Such decisions may be informed by the example of Dutch elm disease (DED) which caused most mature elms in Europe and North America to be lost in the mid-20th Century. Although affected elms can regenerate, they succumb again after several years. Considerable effort has been made to restore elm through: (1) cloning and breeding of surviving local trees, (2) introduction of resistant Asiatic species, (3) introduction or production of hybrids bred from local and Asiatic species. As trees are planted out into the landscape, it is necessary to record them, and monitor their survival. Here, we compile and map 82,244 elm trees in Britain recorded as 269 distinct elm species, hybrids and cultivars. The vast majority of records are for surviving field elm, English elm and wych elm, which would grow again to maturity if the virulence of the pathogen could be attenuated. The most commonly planted elms were the hybrids: ‘Nanguen’ (commercial name Lutece), followed by ‘New Horizon’, ‘Ademuz’, ‘Lobel’ and ‘Sapporo Autumn Gold’, each of which had over 200 occurrences. The current elm-scape of Britain represents much restoration effort and contains considerable genetic diversity.

## Introduction

Elm (*Ulmus* L.) has had a significant role in the UK’s landscape for centuries, as a common broadleaved tree in woodlands and hedgerows with ecological, economic and cultural importance (e.g. Richens 2012; Clouston and Stansfield 1979). However, two global outbreaks of Dutch Elm Disease (DED) during the 20th Century (Brasier and Buck 2001) resulted in a rapid loss of mature elms from the landscape (Potter et al. 2011; Gibbs and Howell 1972, 1974; West et al. 2025). In response, extensive research was initiated including on continuing evolution and biocontrol to attenuate the pathogen (Brasier 2000), and the development of resistant trees, mainly for urban landscape (Martín et al. 2019; Mittempergher and Santini 2004; Holmes and Heybroek 1990; Smalley and Guries 1993). Today, biocontrol has yet to be attempted in the field, but many potentially DED-resistant elm trees have been released and planted in Europe, and to some extent in the United Kingdom and elsewhere (Russell and Buggs 2019). Mapping the distribution of elm species and cultivars is an important step towards monitoring long-term performance of such trees and understanding their suitability for elm restoration projects. Potentially resilient wych elms (*Ulmus glabra* Huds.), derived from controlled crosses of fully mature survivors of disease, have recently been planted in Scotland (Coleman 2025).

The taxonomy of the genus *Ulmus* L. is now underpinned by the molecular and phylogeny of Whittemore et al. (2021), which resolves the ca. 30 elm species known globally into six lineages, assigned to six named sub-generic Sections. The three European elms, Wych elm (*U. glabra*), Field elm (*U. minor*) and White elm (*U. laevis*), each belong to a different Section and thus are only distantly related to each other. Wych elm is the only elm unanimously considered native to the UK, and is most abundant in northern and western regions (Thomas et al. 2018; Clouston and Stansfield 1979; Coleman 2021). In lowland England from Yorkshire southwards, field elm is most frequent. While it might be native, it is often suggested that diverse clones of *U. minor* were introduced to the UK from the Bronze Age for agricultural purposes and some subsequently became widely planted (CABI 2022; Richens 2012). Apart from human influences the differing distributions of wych elm and field elm can be attributed to ecological and climatic preferences of these species, with *U. glabra* able to tolerate lower temperatures than *U. minor*. Northern and western areas of the UK are generally wetter and cooler than the southeast, indicating that *U. minor* is better adapted to the drier conditions of southern England (Coleman et al. 2000).

In Britain, as across Europe, there is high phenotypic variability within *U. minor*, perhaps in part due to its ability to hybridise with *U. glabra* (Stace 2019). Taxonomic treatments have varied from defining *U. glabra* and *U. minor* as a single species (Machon et al. 1995), to a microspecies approach classifying 62 species mainly on leaf morphology (Armstrong and Sell 1996). However, analysis of molecular markers in the *U. minor* microspecies *U. plotii* Druce, endemic to the English Midlands, revealed that 14 samples were the same genotype, indicating that known instances of this ‘species’ endemic to the English Midlands were, in fact, clones (Coleman et al. 2000). The study raised the question of whether such clones should be treated as cultivars, given that elms have a long history of being introduced and planted throughout Britain. Similarly, a study of the genetic sequences of ‘English elm’ (*U. procera* Salisb. or *U. minor* vulgaris), the most abundant ‘field elm’ in England, showed sampled trees to be a single clone probably introduced by the Romans and planted widely in Spain and England (Gil et al. 2004). This cultivar was redesignated *U. minor* ‘Atinia’, after its suggested origin in Atinia, Italy (Gil et al. 2004; Coleman et al. 2016) and will be referred to as such hereafter. Whittemore et al (2021) also provided evidence that some British elm forms ascribed specific epithets are clones of *U. minor,* and in addition confirmed the long-believed hybridisation of *U. minor* and *U. glabra* in Europe. Though various field elm types including English elm may not be from the UK, they have frequently been described as ‘British elms’ (e.g. Shreeve and Seddon 2024). As indicated by the above genetic studies, the impact of human activity on the distribution of elm cannot be ignored. Elm has long been cultivated throughout Europe as a genus of economic and cultural use, probably leading to its introduction into new areas and widespread planting of favoured cultivars.

In the 20th century, two outbreaks of DED have devastated the elm populations of Europe and North America (Peace 1960; Brasier and Buck 2001). Caused by introduced fungal species of the genus *Ophiostoma* and carried by Elm bark beetles (*Scolytus* spp.), the disease results in vascular wilt, defoliation, and death of the tree. *Ophiostoma ulmi,* introduced into Europe and North America in the early 1900s, resulted in a decline of around 30% of mature elms in Britain by 1940 (Brasier and Webber 2019; Peace 1960). The second pandemic was caused by the introduction into both Europe and North America of a much more aggressive species, *Ophiostoma novo-ulmi*, probably as early as the 1940s (Brasier and Kirk 2010; Brasier et al. 2021). Following its subsequent introduction into Britain. *O. novo-ulmi* had caused the loss of most mature elms, amounting to some 25 million by the 1990s (Brasier and Webber 2019). In the current, post - epidemic period a new disease dynamic has emerged in which repeated waves of beetle population expansion occur once elms reach a size large enough to support beetle breeding. This has led to large numbers of small recruitment elms persisting in the landscape but unable to reach full size (Brasier and Webber 2019), though some Wych elms can reproduce sexually before succumbing to DED.

In response to the rapid loss of large elms, efforts to select or breed disease-resistant elms were initiated in Europe and North America. The first elm breeding programme was started in the Netherlands in 1928, led by Christine Buisman (Holmes and Heybroek 1990), initially focussed on identifying native individuals exhibiting disease resistance to *O. ulmi*. This led to the release in 1937 of a clone of *Ulmus minor* named ‘Christine Buisman’ following her death in 1935 (Mittempergher and Santini 2004). Subsequent breeding programmes in the United States, Spain, and Italy and an intensive selection programme in France (Collin et al. 2020; Santini et al. 2008), all involving direct inoculation trials with *O. novo-ulmi,* have led to the production of numerous resistant cultivars and clones mostly aimed at urban or ornamental planting. The introduction of Himalayan (*U. wallichiana*) and east Asian (e.g. *U. pumila*) elms into the crosses has led to increased the resistance of the cultivars available (Martín et al. 2019). Many of these cultivars have been introduced, though in small numbers, into the UK, including some clones that were never commercially released (Fig. 1).

**Figure 1.**
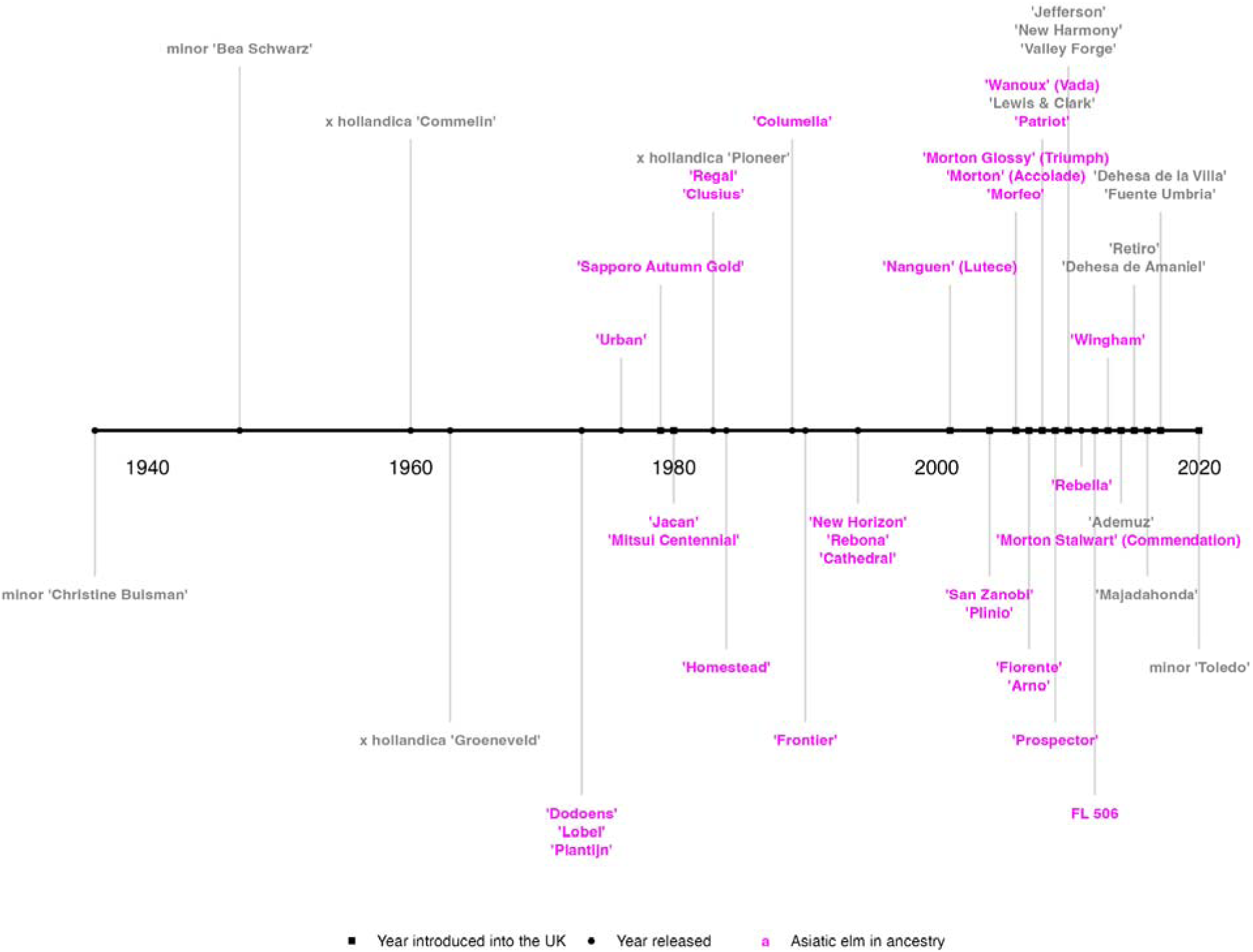
Timeline of disease-resistant elm cultivars, depicting either when they were commercially released or first introduced to the UK (Wikipedia contributors 2024; Mittempergher and Santini 2004). Those with genes from Himalayan or east Asian elms are highlighted.

In Britain, there is great interest in restoration of the enormous visual and cultural contribution of elms in the landscape (Carter et al. 1979; West et al. 2025). In terms of resistant elms, the ideal for British woodlands would probably be forms of or with similar characteristics to, *U. glabra* (Coleman 2025); and for the landscape and hedgerows, a suckering *U. minor* type with fine architectural form, such as English elm, *U. minor* Atinia. As a range of putatively disease-resistant elms are planted in the UK, it is important to monitor their climatic suitability, their growth and form and their survival under high disease pressure (or artificial inoculation) in UK conditions to determine their suitability. Therefore, this study aims to gather elm occurrence data and information from these plantings in order to support long-term monitoring. By improving our understanding of the variety of elm species and subtaxa in the country, trees with the best promising characteristics for restoring British elms can be identified, studied and selected for further research or breeding.

## Methods

### Data Collection

Occurrence records for the genus *Ulmus* L. were accessed from open data repositories in 2024 including the Global Biodiversity Information Facility (Gbif.Org 2024), iNaturalist (https://www.inaturalist.org) and National Biodiversity Network UK atlas (https://nbnatlas.org). Open records of elm trees maintained by councils were also accessed from Greater London Authority, Bristol City Council, Nottingham County Council and Belfast City Council under the Open Government Licence V3.0. In addition, data was received from the Conservation Foundation, Royal Botanic Gardens, Kew, The Royal Parks, Nether Edge and Sharrow Sustainable Transformation, National Memorial Arboretum, Butterfly Conservation, Sir Harold Hillier Gardens, Westonbirt Arboretum, Hampshire Forest Partnership, Hillier Nurseries and Brighton Council’s National Elm Collection and Royal Botanic Garden Edinburgh. Further records were obtained through personal communication with private individuals and elm enthusiasts.

### Data Organisation and Filtering

Records were edited and standardised using OpenRefine and R v4.4.1 using packages dplyr and tidyr (Wickham et al. 2017.). From the names used in each dataset, synonyms and accepted species names were identified using the International Plant Names Index (https://www.ipni.org), Plants of the World Online (https://powo.science.kew.org) and World Flora Online (https://www.worldfloraonline.org). Synonyms for elm cultivars were found using lists of elm synonyms and cultivars on Wikipedia (https://en.wikipedia.org), and Royal Horticultural Society Plant Finder (https://www.rhs.org.uk/plants). Tree status, height, girth, form, planted date and date of recording were included where available. Planting dates were used to calculate ages and sort trees into age groups of 0-15, 16-30, 31+ years.

Records with coordinates outside of the UK or in implausible locations (such as offshore) were excluded. Records without coordinates but with descriptions of location were georeferenced as accurately as possible, or estimated from OS grid reference or postcode. Exact duplicates based on coordinates, name and location description were identified and removed. A distance matrix was produced to group close trees using the R package geosphere (Hijmans 2010). Trees of the same variety within 5m of each other were assumed to be duplicate records and only the most recent observations were used. Shapefiles of Watsonian and Praeger Vice Counties were accessed from github (https://github.com/BiologicalRecordsCentre/vice-counties; https://github.com/SK53/Irish-Vice-Counties) before merging and clipping to the extent of the UK using QGIS software (https://qgis.org). The sf package (Pebesma 2016) in R was then used to assign country and vice-county information to each data point.

### Distribution Mapping

Base maps of the UK were obtained as shapefiles from GADM v4.1. The filtered data was used to plot point distribution maps for each elm species or cultivar using the R packages sf (Pebesma 2016) and ggplot2 (Wickham 2016). In cases of species with high point density, such as *U. glabr*a and *U. minor* ‘Atinia’, the 20km OS grid (https://github.com/OrdnanceSurvey/OS-British-National-Grids) was applied to visualise the abundance of records per grid square.

### Database

The collated records were compiled into a .csv file, including columns “Source”, “Verbatim name”, “Name”, “Tree Status”, “Tree Parent”, “Latitude”, “Longitude”, “OS Grid Reference”, “Location Description”, “Access”, “Tree Description”, “Height (m)”, “Girth (m)”, “Planted Year”, “Age”, “Veteran status”, “Date recorded”, “Year Recorded”, “url”, “Licence”, and “Rights Holder”. Some records were omitted from the database to protect privacy.

### Collating Records from Conservation Foundation Campaigns

In the 1980s the Conservation Foundation (https://conservationfoundation.co.uk) participated in the “Elms Across Europe” campaign (https://conservationfoundation.co.uk/history accessed 15/04/2026) and distributed ‘Sapporo Autumn Gold’ trees to UK schools. A list of 250 participating schools was collated from their archives. To update records, schools that are still operating at the same address were contacted to request information on the tree’s status.

The Conservation Foundation’s Great British Elm Experiment launched in 2009 (https://conservationfoundation.co.uk/history accessed 15/04/2026) and aimed to plant cuttings of surviving mature trees, in an effort to trial their resistance (Shreeve and Seddon 2024). Fourteen parent trees of British elms (including *glabra*, *minor*, *minor* ‘Atinia and *x hollandica*) were propagated and planted throughout the UK for monitoring as a citizen science project. Over 3,000 cuttings were distributed to participating schools, community groups and individuals, who could then register their elm and provide updates on the Great British Elm Experiment map (https://conservationfoundation.co.uk/elm-experiment/ accessed 23/1/2026). We collated these records including updates regarding their location and alive/dead status within 5 years of planting.

## Results

In total, 125,197 records of elm occurrences were obtained, including human observations, preserved specimens and living specimens in botanic gardens and arboreta. After filtering for duplicates, this number decreased to a total of 83,275 distinct records with 82,244 including sufficient location information for mapping.

### Taxonomic representation

After assigning synonyms to some names, the data set represented 35 species, 3 subspecies, 13 varieties, 1 form, 39 hybrids and 178 cultivars or clones (Fig. 2), in the opinion of the people recording the occurrences. Of the 178 cultivars, 73 were cultivars or clones of disease-resistant elms. Records for 13 hybrids, 2 varieties and 3 cultivars could not be georeferenced and were therefore not mapped.

**Figure 2.**
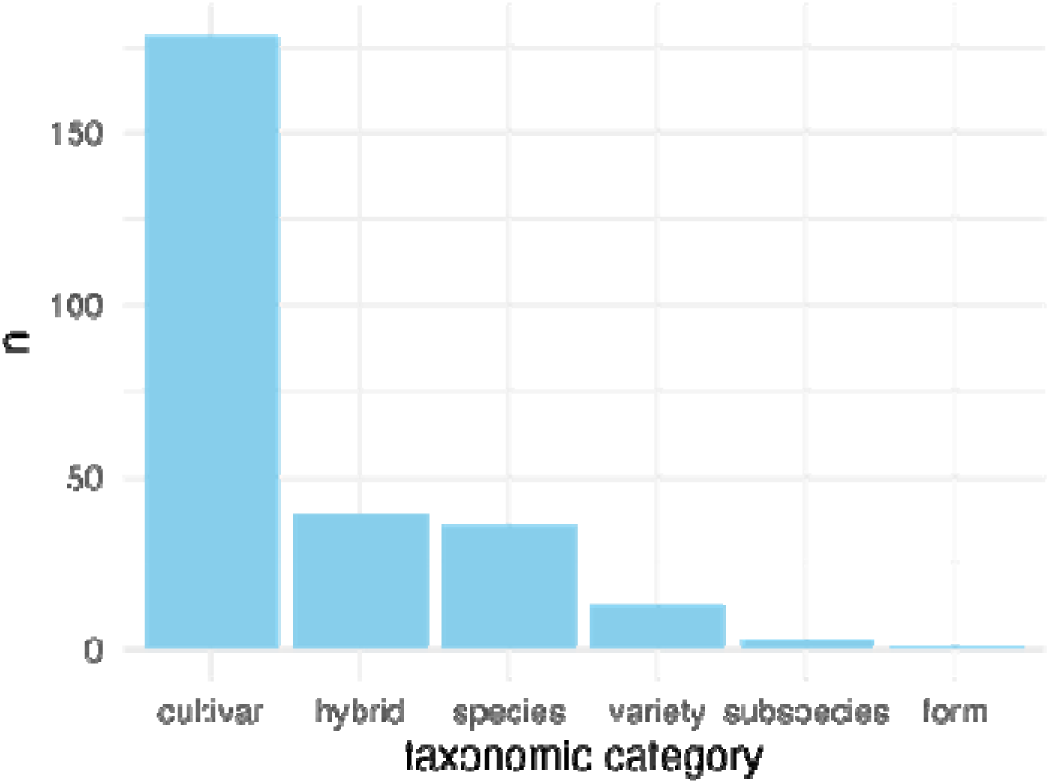
Number of distinct names per taxonomic category in the database.

It was not possible to determine the correct synonym for four trees identified as *Ulmus japonica*, as these could represent two different taxa depending on the species author. *U. japonica* Siebold is an accepted synonym for *U. parvifolia*, and *U. japonica* Sarg a synonym for *U. davidiana var. japonica*. These records were therefore kept as *U. japonica* in the dataset, but this species name has not been included in the total number of species. No species name was listed for 12.6% of records; these were identified only to the genus level (*Ulmus* spp.), demonstrating the uncertainty of identifying some elms.

### Temporal Distribution of Records

As 94.8% of the records included information of the year recorded, the temporal distribution of these occurrences was examined. The occurrence records covered a period of over 200 years, however, a large increase in the number of records could be seen in the late 20th Century (Fig. 3.). Elm records since the year 2000 represent over 53% of the dataset, with an average of 1,774 records per year between 2000 and 2024. Even when filtered for the most recently recorded trees, the dataset included 240 elm taxa and subtaxa.

**Figure 3.**
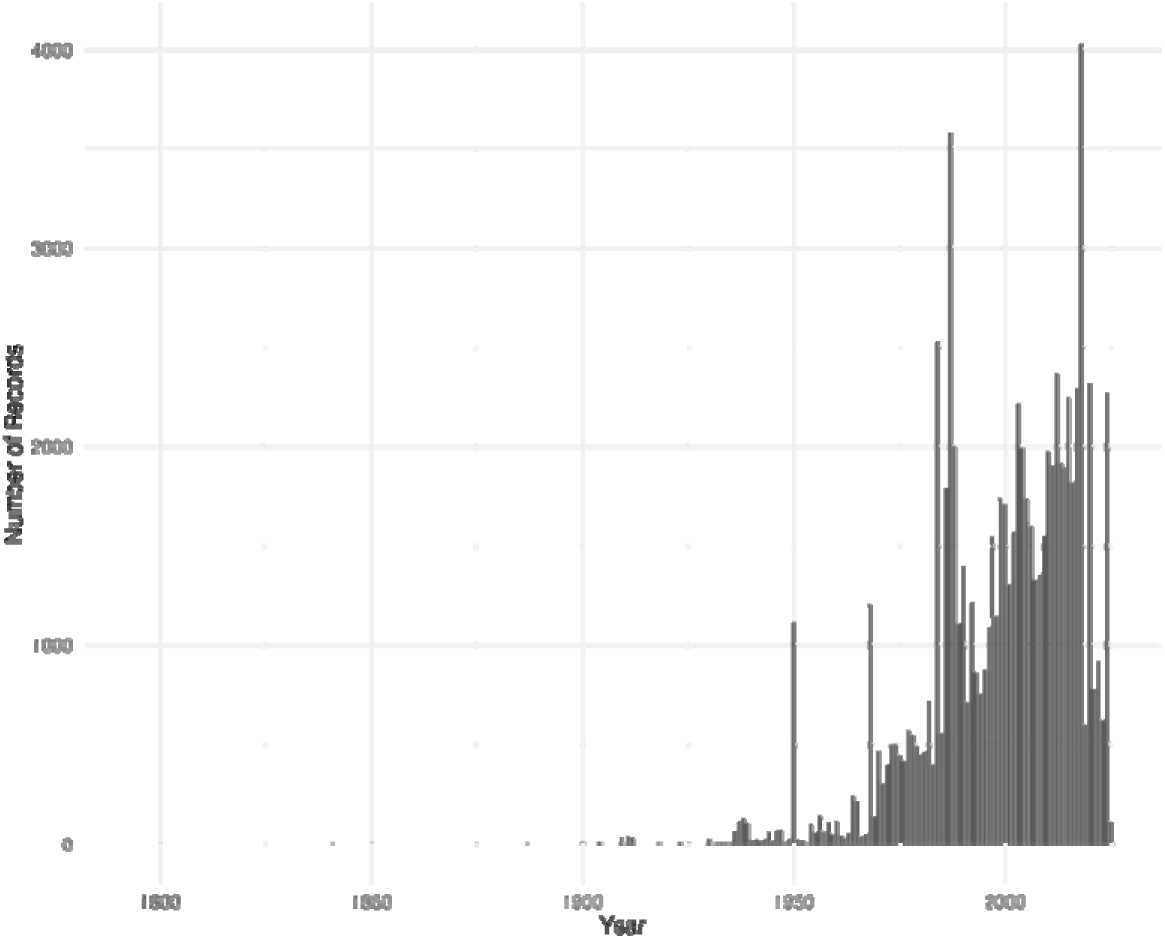
Number of elm occurrences recorded per year in the database

### Distribution of Ulmus glabra, U. minor and U. minor ‘Atinia’

Point distribution maps of *Ulmus glabra*, *U. minor* and *U. minor* ‘Atinia’ showed a large number of occurrences throughout the UK, with *U. glabra* more commonly recorded in northern and western regions compared to *U. minor* (Fig. 4a, c, e). Maps showing frequency of records per 20km OS grid indicate areas of high point density (Fig. 4b, d, f). Similar distribution patterns were seen when mapping just the records obtained from the year 2000 onwards (Fig. 5.).

**Figure 4.**
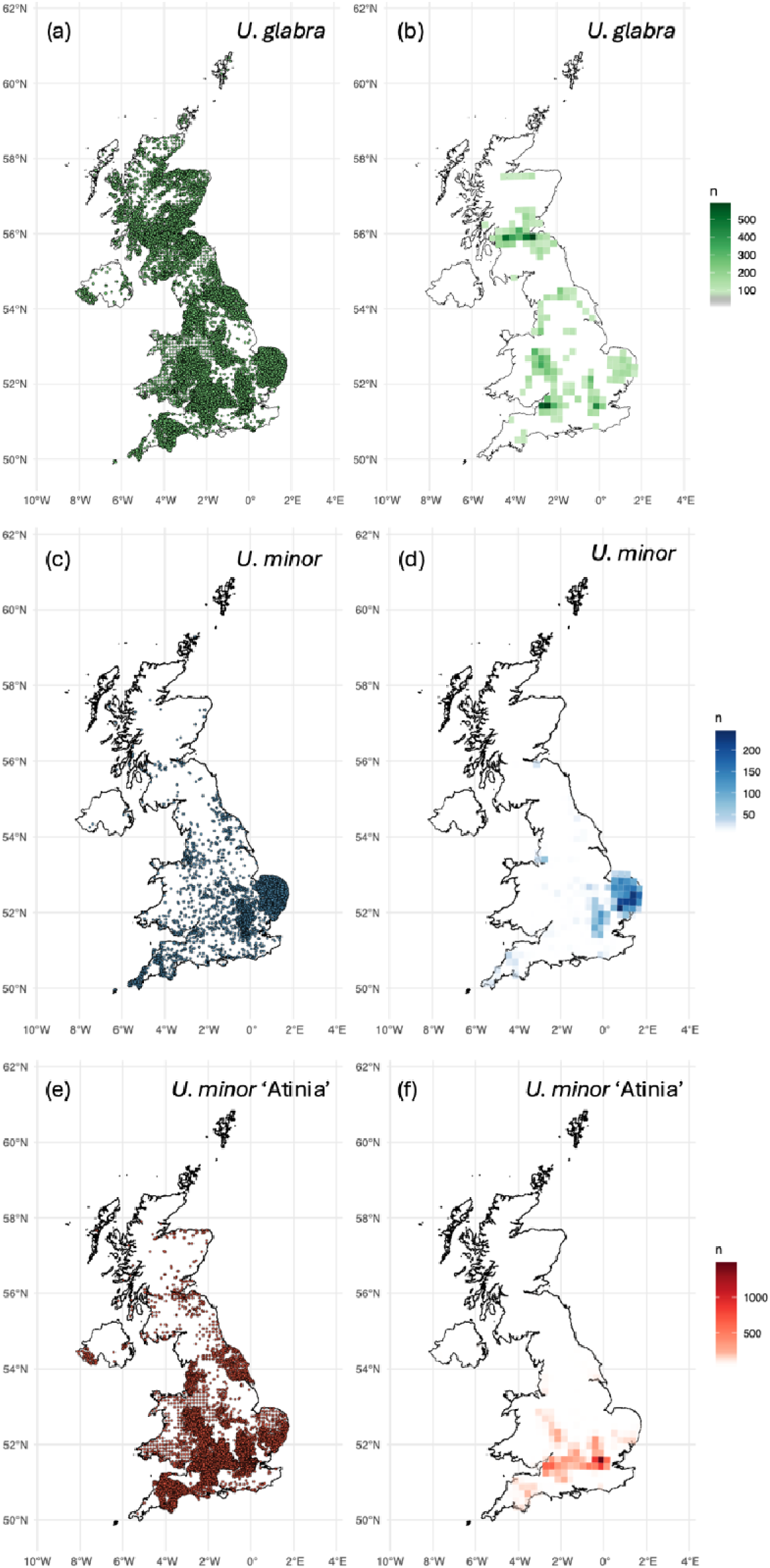
Distribution maps of *Ulmus glabra, Ulmus minor* and *Ulmus minor* ‘Atinia’ recorded in the database.

**Figure 5.**
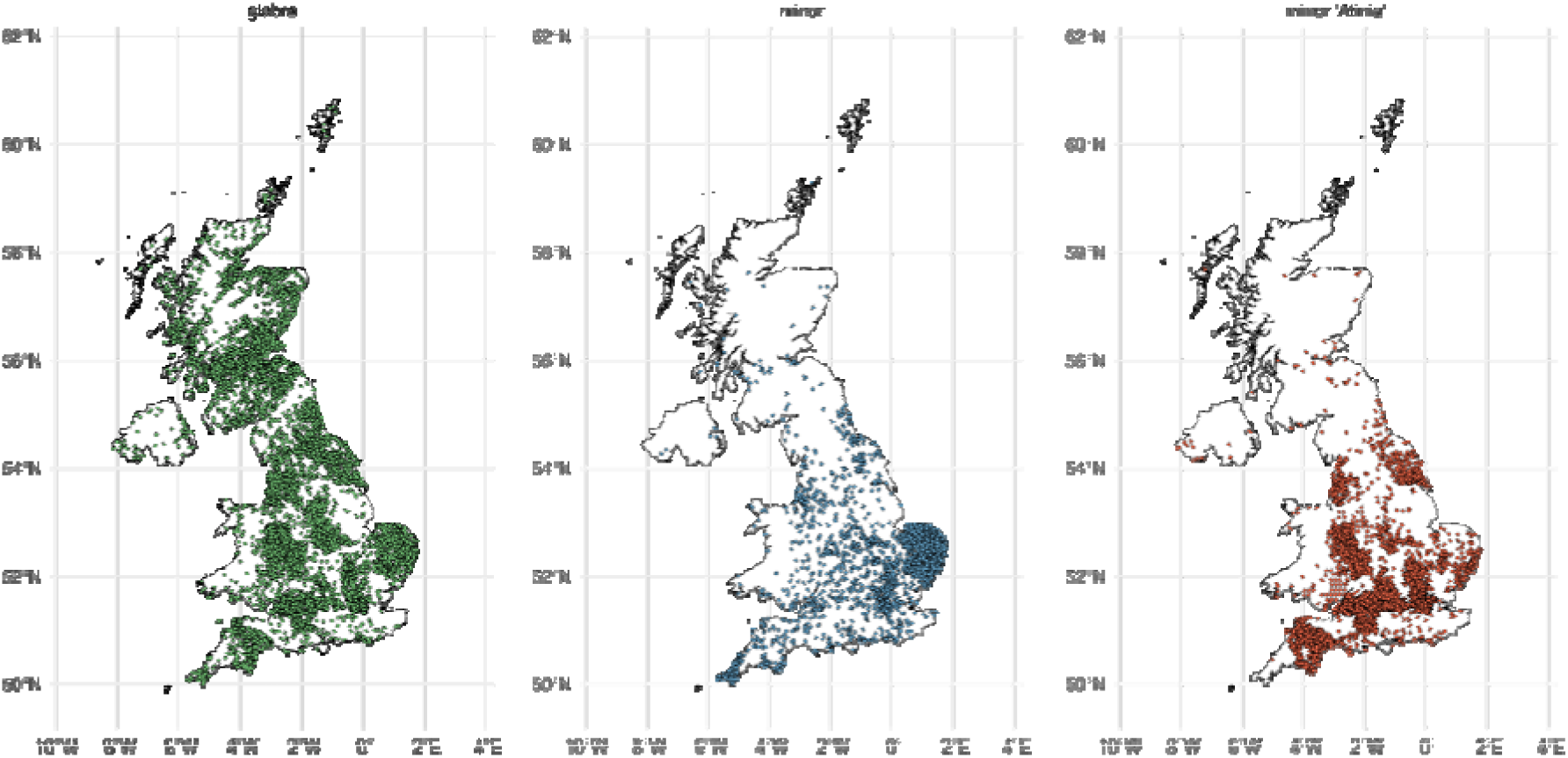
Point distribution maps of records of *U. glabra*, *U. minor* and *U. minor* ‘Atinia’ occurrences recorded after the year 2000.

### Age distribution of surviving trees

It was possible to calculate the ages of 4,228 trees from the data gathered. The majority (79.8%) of these trees were described as between 0-15 years of age at the time of recording (Table 1), which is primarily due to the availability of data from recent planting projects. In addition to the 326 elms known to be over 30 years old, 490 elms were recorded as a ‘Veteran’ or ‘Ancient’ tree, and a further 148 were noted as mature in descriptions (Fig. 6a). This includes records of surviving ‘British elms’ (Fig. 6b) as well as 114 occurrences of mature disease-resistant cultivars.

**Figure 6.**
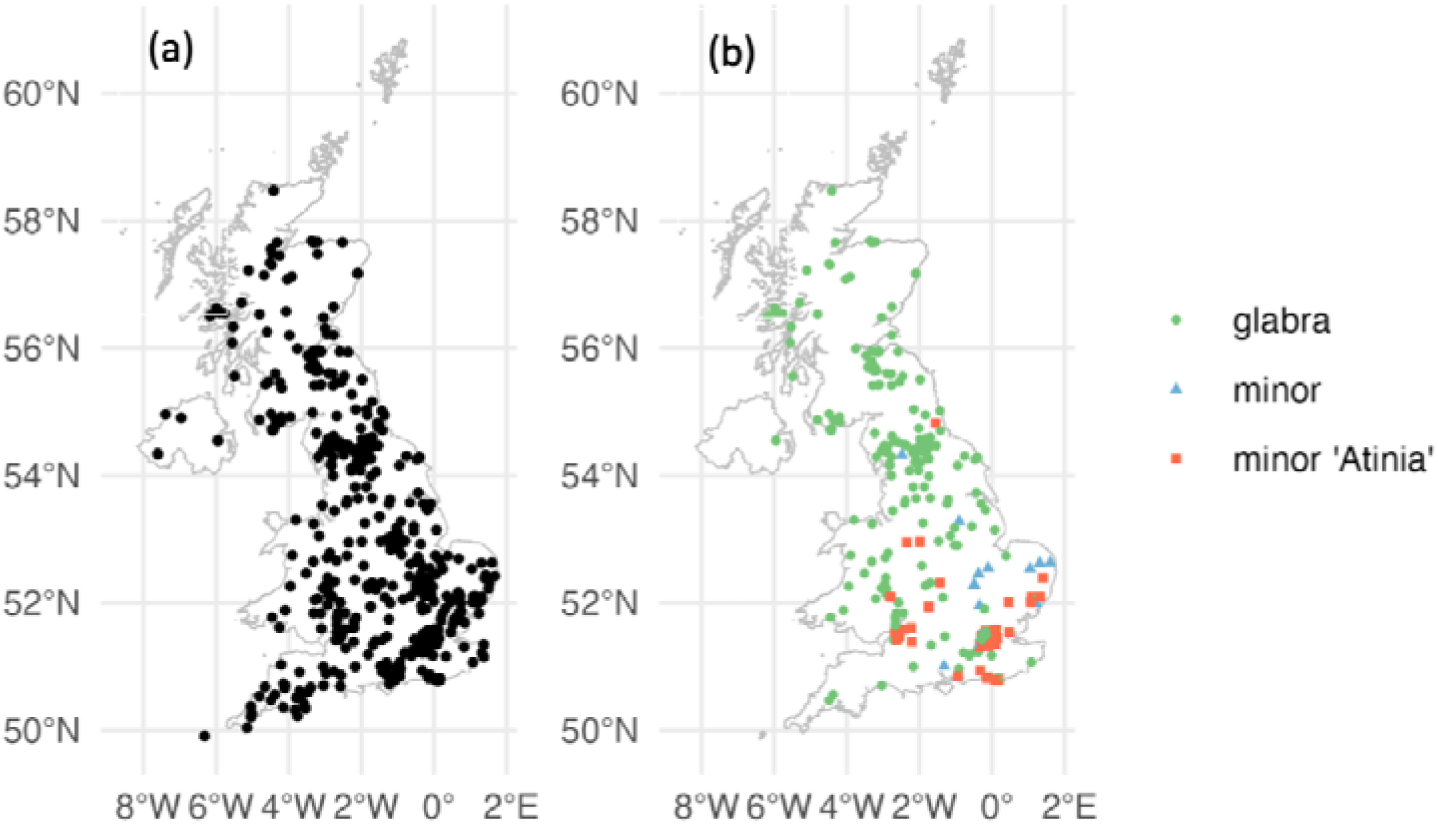
Distribution of a) all 964 records of trees over 30 years old or described as veteran/mature. b) Mature *Ulmus glabra*, *Ulmus minor* and *Ulmus minor* ‘Atinia’ specimens

**Table 1.** Number of elm records within each age group.

| Age Group | Number of records |
| --- | --- |
| 0-15 years old | 3350 (79.8%) |
| 16-30 years old | 519 (12.4%) |
| 31+ years old | 326 (7.8%) |

### Distribution of introduced species and their cultivars

The most widespread introduced species in the UK is the European White Elm, *Ulmus laevis,* with 303 mapped records (Fig. 7b). Other non-native species that have been planted in the UK include North American species such as *U. americana* and *U. rubra*, and Asiatic species such as *U. pumila*, *U. parvifolia* and *U. wallichiana*. These species and their cultivars are more sporadic in their distribution, which is indicative of their use as ornamental trees (Fig. 7a,c,d,e,f).

**Figure 7.**
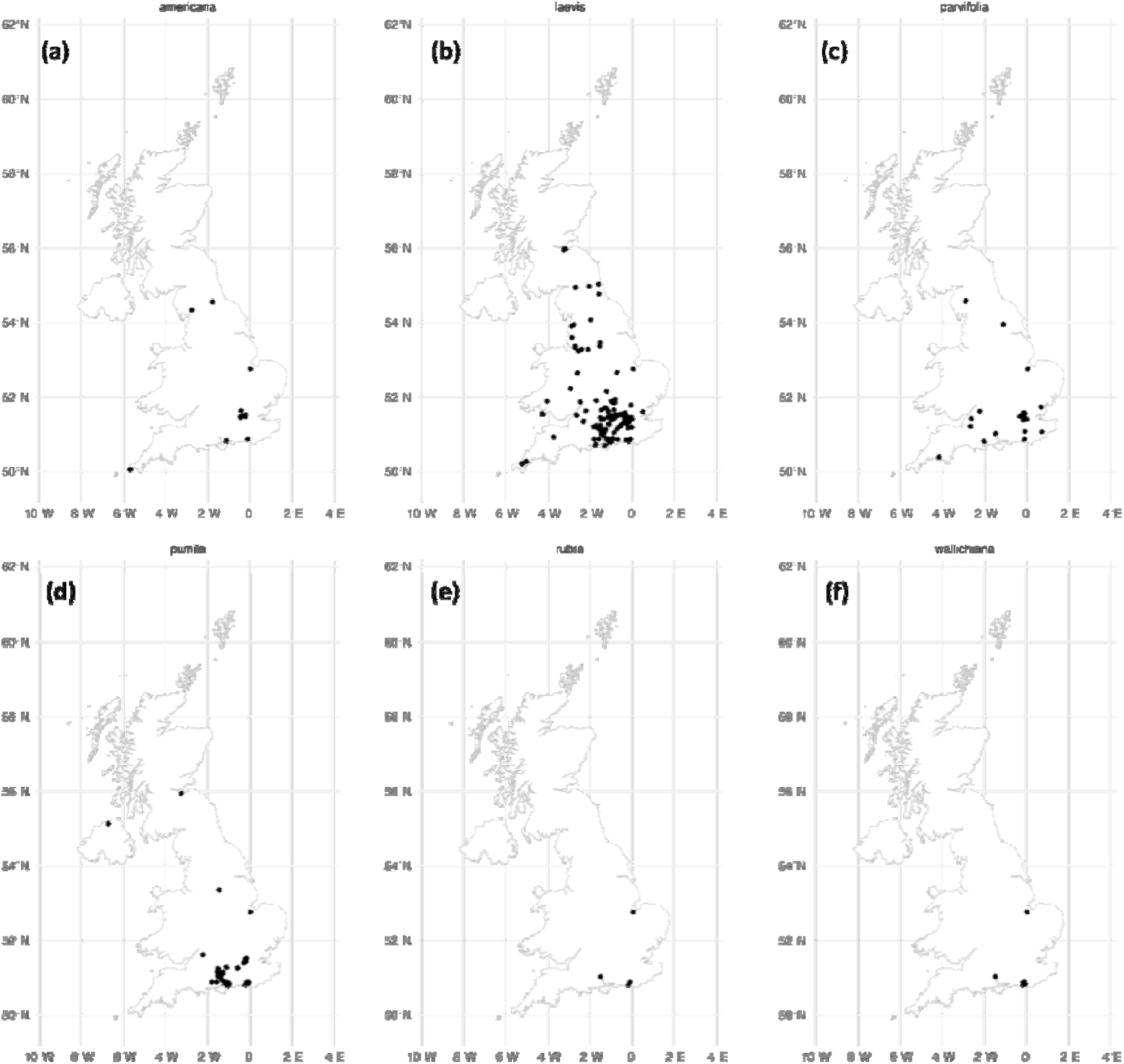
Distribution of records of introduced elm species a) *U. americana*, b) *U. laevis*, c) *U. parvifolia*, d) *U. pumila*, e) *U. rubra* and f) *U. wallichiana*

### Distribution of disease-resistant cultivars

A total of 4,757 records for disease-resistant elm cultivars were collated in the dataset, representing 72 different cultivars or clones. These records covered 57 out of the 122 Watsonian and Praeger Vice Counties of the UK, with the highest number of records in South Hampshire, North Hampshire and Middlesex. Of the 72 cultivars introduced to the UK, *Ulmus* ‘Nanguen’ (commercial name Lutece) had the highest number of occurrences followed by ‘New Horizon’, ‘Ademuz’, ‘Lobel’ and ‘Sapporo Autumn Gold’ (Fig. 8).

**Figure 8.**
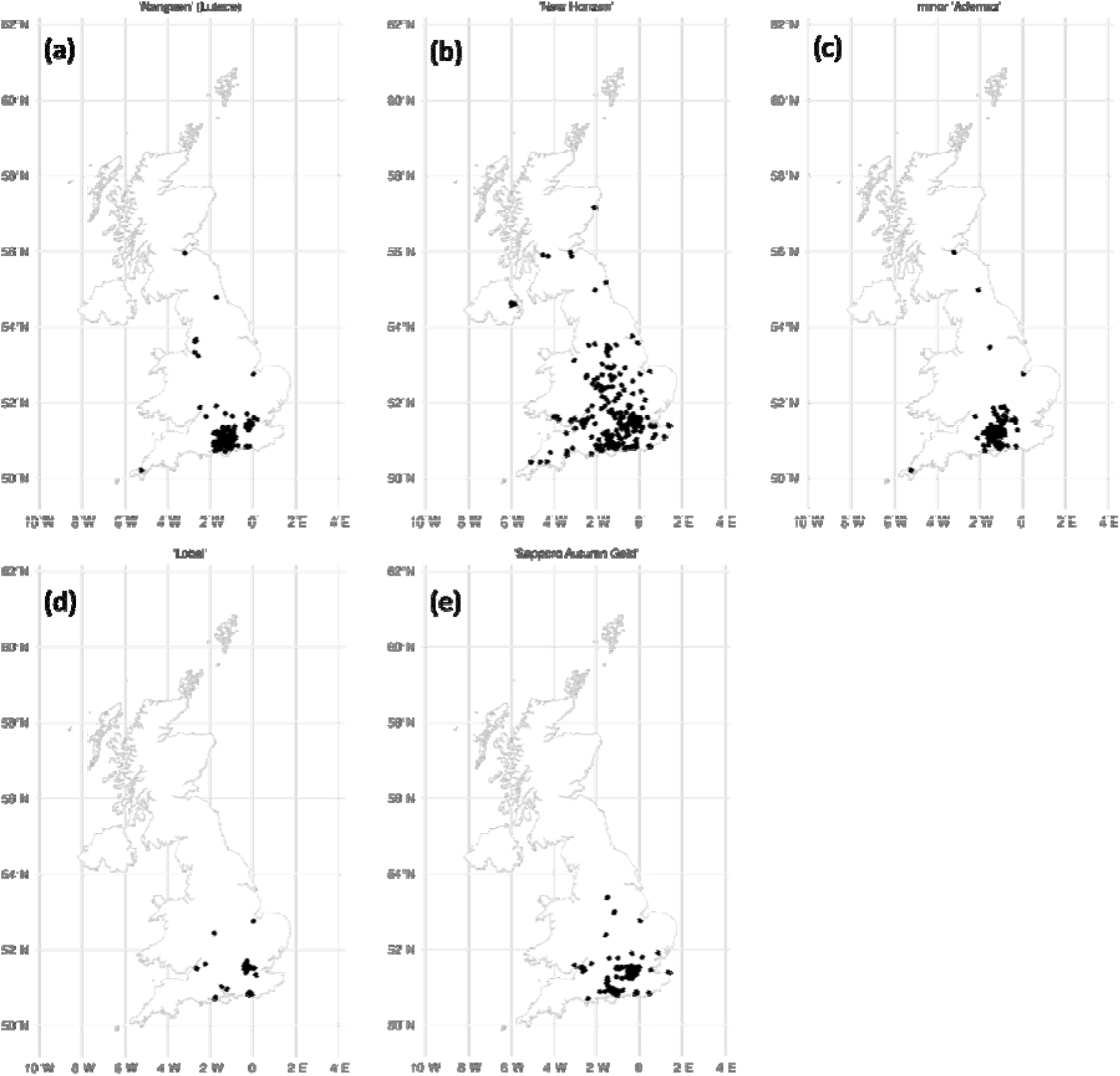
Distribution of records of the five most frequently recorded disease-resistant elms a) ‘Nanguen’ (Lutece), b) ‘New Horizon’, c) ‘Ademuz’, d) ‘Lobel and e) ‘Sapporo Autumn Gold’

Twenty-three resistant cultivars were rare in occurrence with ≤ 2 records, located in Royal Botanic Gardens, Kew, Grange Farm Arboretum, Sir Harold Hillier Gardens or the National Elm Collection in Brighton. This includes several clones from the Italian breeding programme that were not commercially released (FL 214, FL 316, FL 441, FL 465, FL 626, FL 610).

### Collating Records from Conservation Foundation Campaigns

It was possible to contact 137 of the 250 schools listed as participating in the 1980s ‘Elms Across Europe’ campaign (Fig. 9a). Responses were received from 28, and only 4 schools had knowledge or memory of this elm on their site. Out of these 4 trees, 3 were reported to be alive and healthy, and 1 had been removed in 2007 due to disease (Fig. 9b). The four confirmed locations of ‘Sapporo Autumn Gold’ were then added to the dataset.

**Figure 9.**
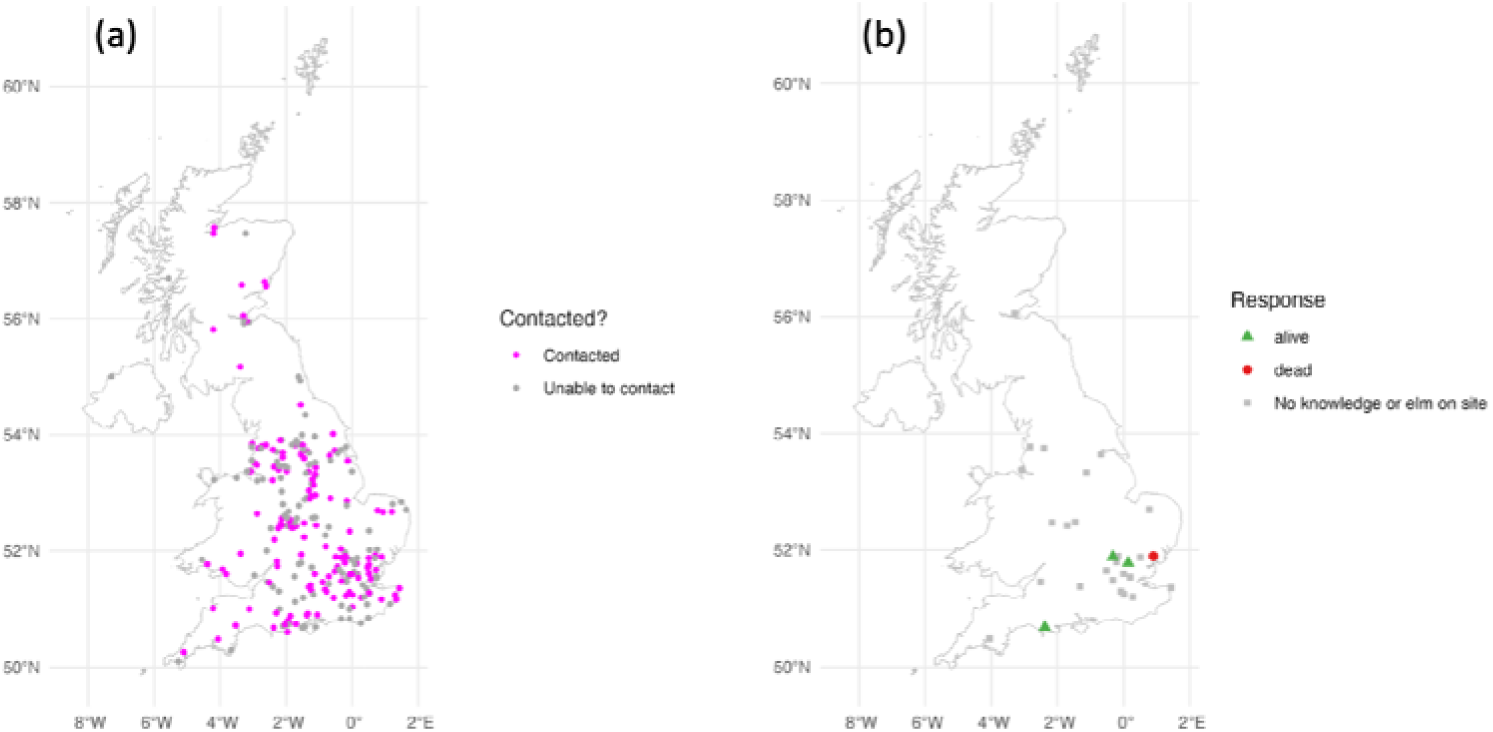
a) Map of schools listed as participating in the ‘Elms Across Europe’ campaign, those that were contacted are highlighted in pink. b) Map of the responses received

We collated 1,279 records from The Conservation Foundation’s Great British Elm Experiment launched in 2009 (Shreeve and Seddon 2024), and records of their survival after 5 years (Table 2, Fig. 10.). To see if these trees had potentially been recorded more recently through other sources, we also compared the locations of these trees to the groups produced by the distance matrix; however, no overlapping records were found.

**Figure 10.**
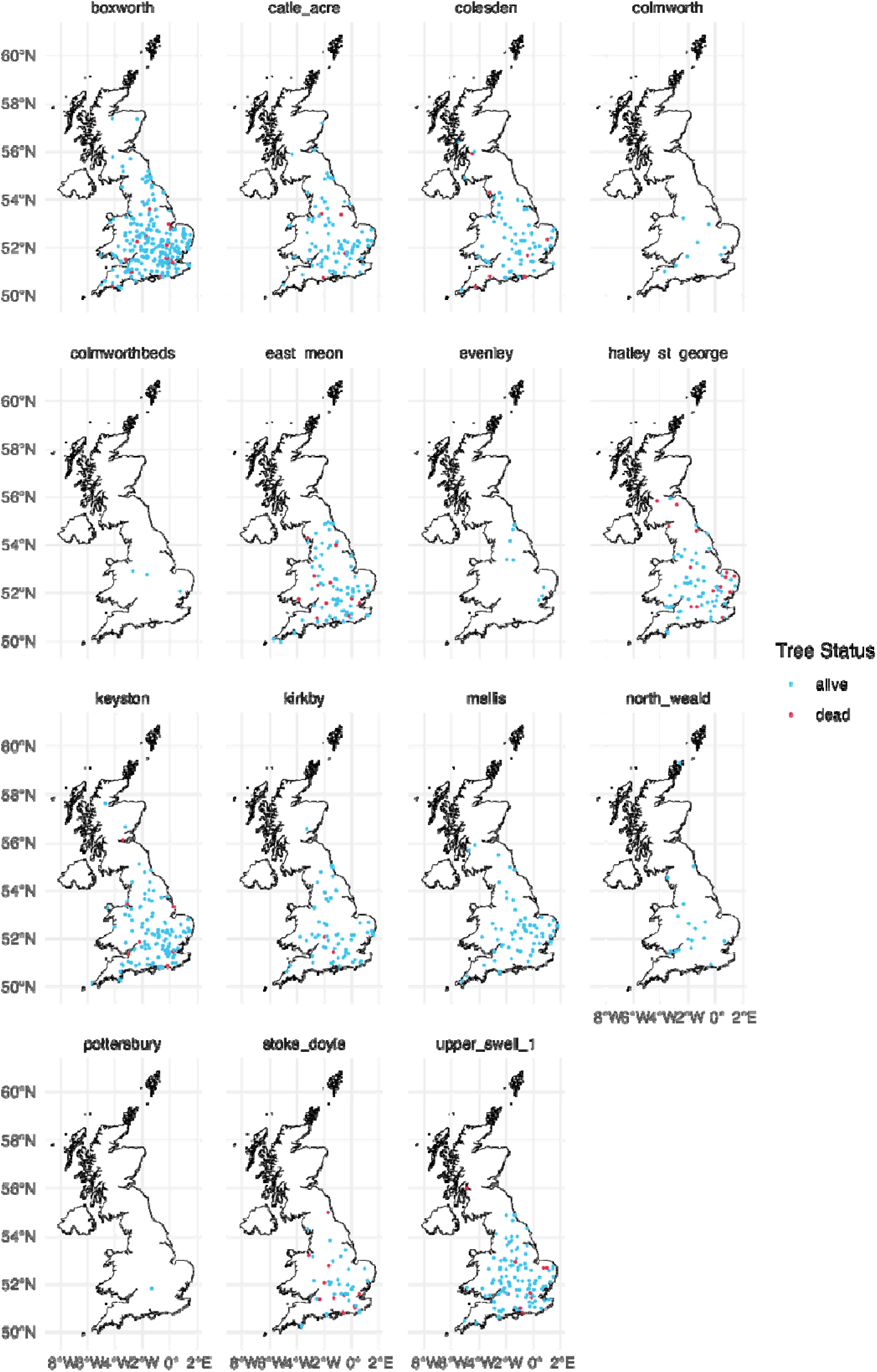
Distribution of elms planted in the Great British Elm Experiment by tree parent. Elms reported to have died within the first five years are shown in red and those alive in blue

**Table 2.** Number of elms reported as alive or dead (within 5 years of planting) per parent tree of the Great British Elm Experiment.

| **Tree Parent** | **Species** | **Alive 2010-2014** | **Dead 2010-2014** |
| --- | --- | --- | --- |
| Boxworth, Cambs | *U. minor* | 303 | 26 |
| Castle Acre, Norfolk | *U. x hollandica* ‘Vegeta’ | 100 | 4 |
| Colesdon, Beds | *U. minor* | 74 | 8 |
| Colmworth, Beds | *U. minor* | 15 | 0 |
| East Meon, Hants | *U. glabra* | 81 | 14 |
| Evenley, Northants/Bucks | *U. minor* | 10 | 0 |
| Hately St George, Cambs | *U. minor* | 74 | 17 |
| Keyston, Cambs | *U. minor* | 131 | 8 |
| Kirkby Stephen, Cumbria | *U. glabra* | 85 | 5 |
| Mellis, Suffolk | *U. minor* | 88 | 1 |
| North Weald, Essex | *U. minor* | 21 | 0 |
| Pottersbury, Northants | *U. minor* | 3 | 0 |
| Stoke Doyle, Northants | *U. minor* | 45 | 16 |
| Upper Swell #1,<br>Gloucestershire | *U. minor* | 139 | 9 |

## Discussion

The data collated here agree with the many previous surveys and descriptions since ca 1962 indicating that *U. minor* (including much *U. minor* ‘Atinia’) is frequent throughout southern England and East Anglia, and *U. glabra* more frequent in northern Britain (Thomas et al. 2018; Coleman et al. 2000; Clouston and Stansfield 1979). The records we collated of *U. glabra* in central and southern England are likely to contain a high proportion of misidentifications as sucker growth of *U. minor* and its hybrids is easily mistaken for wych elm (Coleman 2021).

The increase in the number of records we found from the late 20th Century may be partly due to the increase in biological recording efforts across the country with the establishment of the Biological Records Centre in 1964 (Roy et al. 2015). The large number of elm records since the year 2000, especially of planted exotic species and hybrids with putative or tested resistance to DED, may reflect renewed interest in elm and increased rates of planting of diverse elm taxa.

By collating information from disparate sources, and mapping the locations of *Ulmus* in the UK, this project can support future monitoring and research of DED resistance. The need for such detailed information is demonstrated by the difficulty we found in tracking down the ‘Sapporo Autumn Gold’ elms planted in the 1980s, many of which had been forgotten. Documenting the locations of cultivar trials and elm planting projects as accurately as possible can enable knowledge gaps on their long-term growth and suitability to be filled.

Several recent elm restoration projects have mainly focused on outplanting the offspring of surviving ‘native’ British elms. The Conservation Foundation’s Great British Elm Experiment (Shreeve and Seddon 2024), and the work of King & Co Tree Nursery (https://kingco.co.uk/trees/elm-trees-ulmus-species accessed 23/1/2026) have focused mainly on the planting of clones of large surviving *U. minor trees*. The Royal Botanic Garden Edinburgh’s Scottish Plant Recovery project (Coleman 2025) focuses on *U. glabra* trees surviving in areas where Dutch Elm Disease is prevalent and generates seed from controlled pollinations between survivors. Such projects do not tend to involve testing for resistance by artificial inoculation with *O. novo-ulmi (Pinon et al. 2005)* and some may be ‘disease escapes’ by virtue of ‘field tolerance’ or of low disease pressure. Thus long-term monitoring is essential to understand the level of resistance in these trees.

Although *U. laevis* is the most common non-native elm, it is highly susceptible to *O. novo-ulmi (Brasier and Gibbs 1976)* but less fed on by the *Scolytus* bark beetles (Webber 2000), probably because the presence of alnulin in its bark makes it less appealing to the beetles (Martín-Benito et al. 2005), reducing its likelihood of becoming infected. This ‘field resistance’ trait has led to the incorporation of *U. laevis* in resistance trials (Brookes 2020) and planting projects, with 289 out of the 428 *U. laevis* records from trees known to have been planted in the last 10 years.Recently some recent U. laevis plantings have begun to suffer significant mortality under high disease pressure (pers obs.)

Some resistant elm cultivars have been planted in much larger numbers than others, such as ‘Nanguen’ (Lutece). This could be due to availability from suppliers, as some clones were never commercially released and many are protected by plant breeders rights. However, by focusing planting efforts on a small range of cultivars with the same genetic background, the resilience of these elm stands may be at risk if other elm diseases arrive in Britain, such as Elm yellows (*Candidatus Phytoplasma ulmi*) (Martín et al. 2019; Zhao and Wei 2022) or *O. himal-ulmi* (Brasier 1995).

Not all conservationists are willing to plant elms that contain genetic material from non-native species in all settings. Of the five most frequently recorded disease-resistant elm (Fig. 9), all are hybrids and two have no European native component. ‘New Horizon’ and ‘Sapporo Autumn Gold’ are F_1_ hybrids of two Asiatic species (*U. davidiana* var. *japonica* × *U. pumila*). Lutece and Lobel are complex hybrids containing *U. minor*, *U. glabra*, and *U. wallichiana* and *U. minor*, and Lobel additionally contains *U. carpinifolia* material. ‘Ademuz’ is a selection from Spain, and appears to be a backcross hybrid between *U. minor* and *U. glabra (Vatanparast et al. 2026)*. Thus, such trees may not be considered suitable for some conservation sites, such as Sites of Special Scientific Interest.

The planting of exotic elms can lead to spread of their genetic material. In the landscape of Spain and North America genetic introgression has occurred between planted *Ulmus pumila* and local elm species (Martín et al. 2019). This may have the benefit of spreading resistance, but if conservation goals are to protect and conserve native germplasm, monitoring the spread and pattern of hybridisation between introduced and native elms could be necessary. Monitoring the growth of disease-resistant cultivars in different locations will also improve knowledge of how environmental conditions impact their long-term survival. For example, it is known that environmental factors, such as temperature and light intensity (Sutherland et al. 1997) and soil moisture content, increase or decrease disease resistance in elms (Martín et al. 2019). This is particularly important in the context of climate change, so that “climate-smart” elm restoration efforts can be implemented.

The disease-resistant cultivars currently available are often criticised for failing to replicate the distinctive form of the iconic Atinia clone (Brookes 2020). Britain contains elm genetic material for DED resistance and for the Atinia form, but these have not yet been successfully combined. Future breeding programmes may be able to accomplish this. Though the form of resistant elms differ, many still hold ornamental value which is demonstrated by their distributions in urban environments and parks.

It is important to note that the data gathered in this project is not exhaustive, and the distribution maps should be treated as representative rather than exact. There are also likely to be inaccuracies in the dataset, as elms are notoriously prone to misidentification and some locations were estimated or obscured. As observations often describe multiple trees in one record, the number of records cannot be used to determine population sizes. Anecdotally, we currently see increasing interest in elms, and increased planting activity, so our results here for planted trees will soon be out of date.

### Concluding remarks

As the number of tree epidemics increases, elm provides an important example of what can be expected to happen in the wake of a devastating and ongoing epidemic. The current elmscape of Britain continues to be dominated by short-lived trees that have regenerated from pre-epidemic trees. There is also significant planting of increasing numbers of exotic elm species, hybrids and cultivars that show more resistance to Dutch elm disease. These represent sustained human effort to recover the elmscape by the addition of trees that can grow to maturity. Monitoring is essential to understand the success of these efforts. The future elmscape of Britain will depend on continued human intervention, including new technologies. Understanding human motivations and attitudes to new technologies is key to predicting this future (West et al. 2025). The breeding or genetic engineering of an elm with the iconic form of the Atinian elm and resistance to DED would be a major breakthrough. A reduction in the cost of existing DED-resistant trees, and increased confidence in their long term growth and form in British conditions, could also lead to more planting. However, we should not forget that if the DED were to be attenuated either by biocontrol or a natural attenuation of the pathogen, thousands of existing young trees could survive to maturity and restore the elmscape by natural means. On the other hand, evolution of greater virulence in the pathogen, or introduction of new pathogens, could make the situation much worse.

## Data availability

The database of elms compiled in this study is available at https://doi.org/10.5281/zenodo.15131299

## Author contributions

Catherine Walter compiled and analyses the data and wrote the first draft of the manuscript, Mohammad Vatanparast supervised the compilation and analysis of the paper, Camilla Quintero-Berns compiled and analysed data, David Shreeve provided data and advice, Joan F. Webber provided samples and advice and Clive Brasier contributed to the manuscript, Richard Buggs obtained funding, managed the project and led the final draft of the manuscript.

## Acknowledgements

This work was funded by the Department for Environment, Food & Rural Affairs (Defra) through the Centre for Forest Protection (Ref. 2967). Catherine Walter and Camila Quintero-Berns were supported by a Centre For Forest Protection internship funded by Defra. Richard Buggs and Mohammad Vatanparast were funded by the Centre for Forest Protection project (Ref. 2967). We thank Max Coleman for helpful conversations and comments on an earlier draft of this paper.

## Notes

https://doi.org/10.5281/zenodo.15131299

### Competing Interest Statement

The authors have declared no competing interest.

