## Supplementary Figures for "Mapping the distribution of elms in Britain after a century of Dutch elm disease"

### Maps for all 253 elm species and cultivars with location information

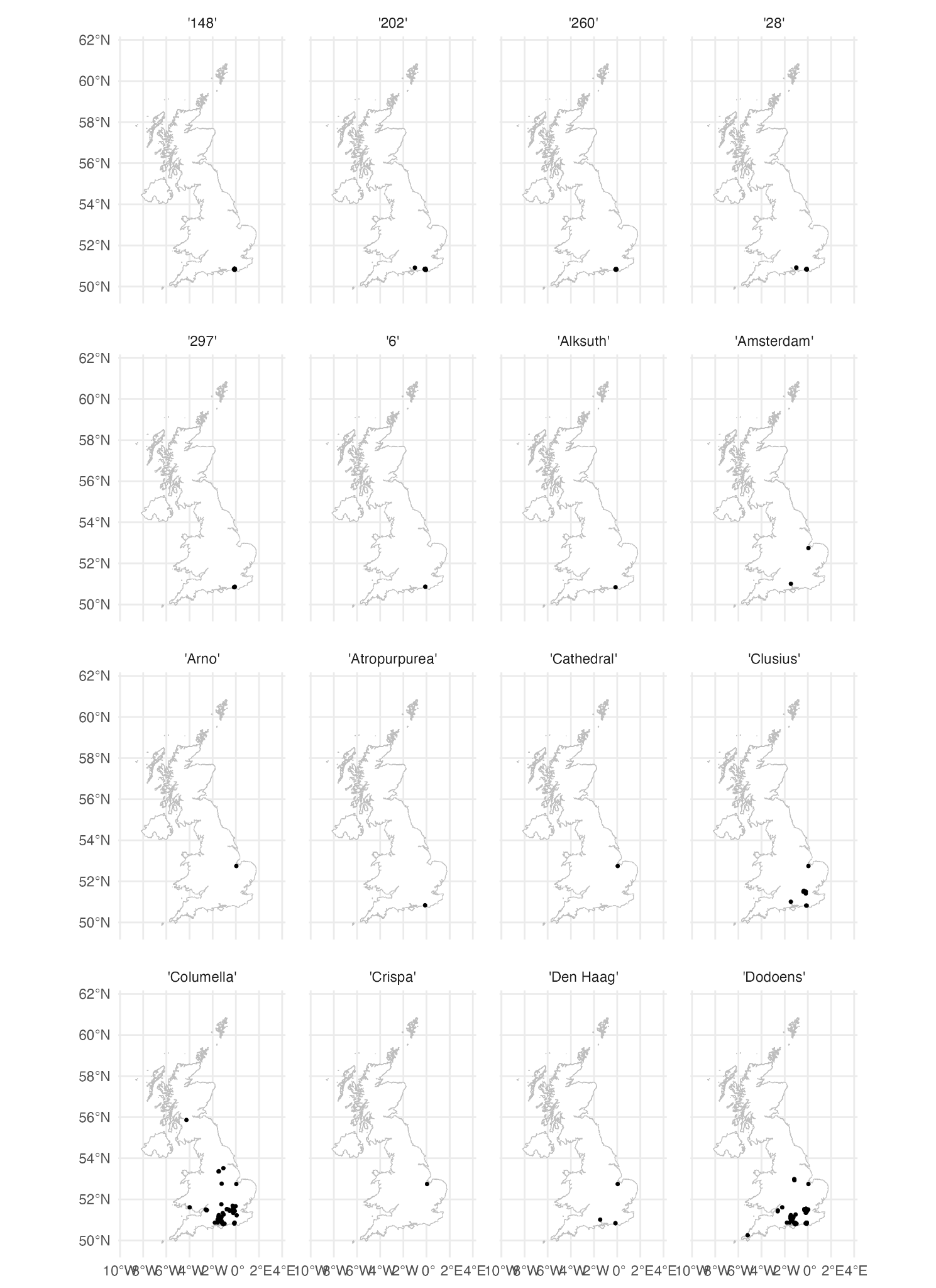

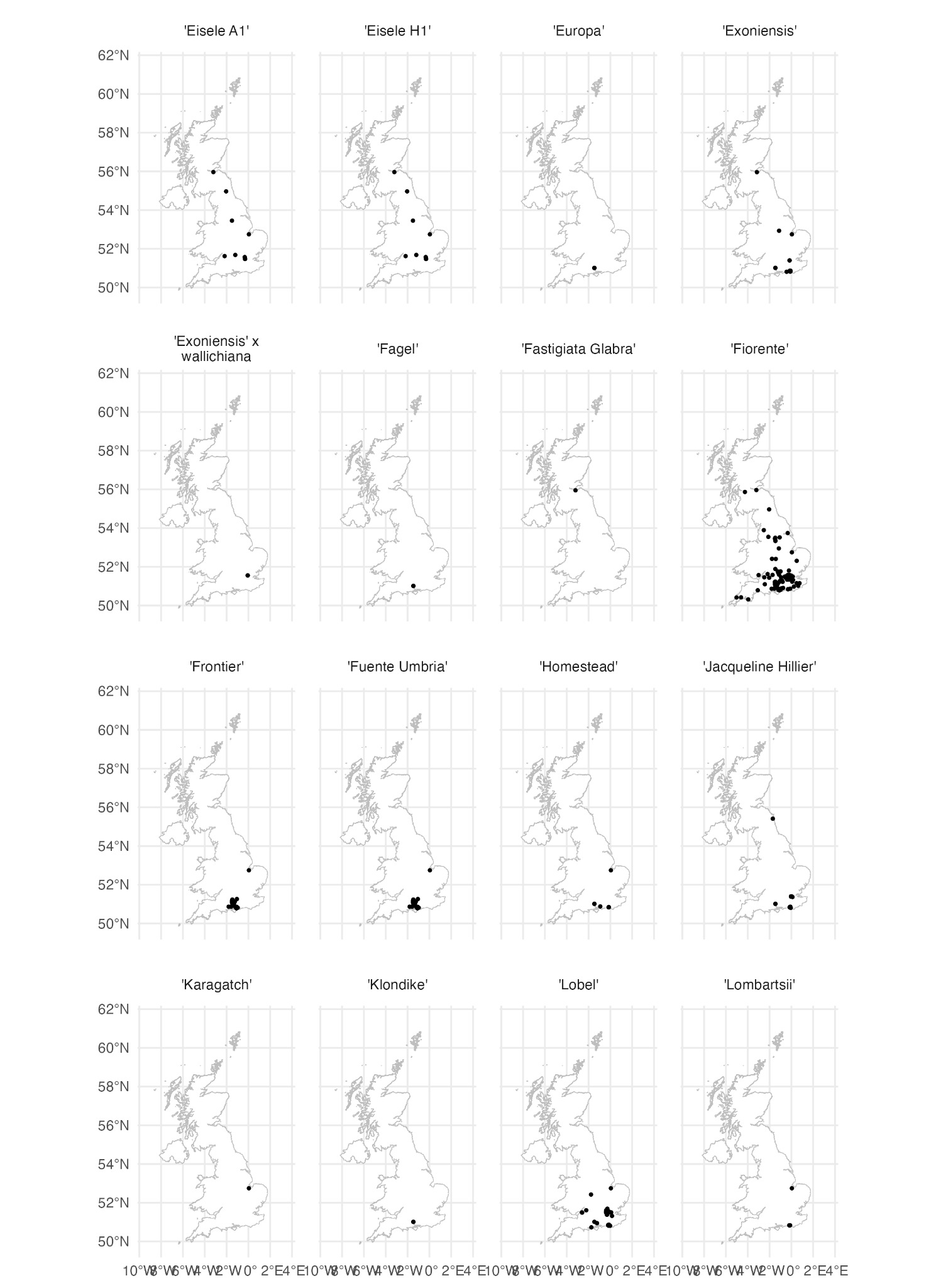

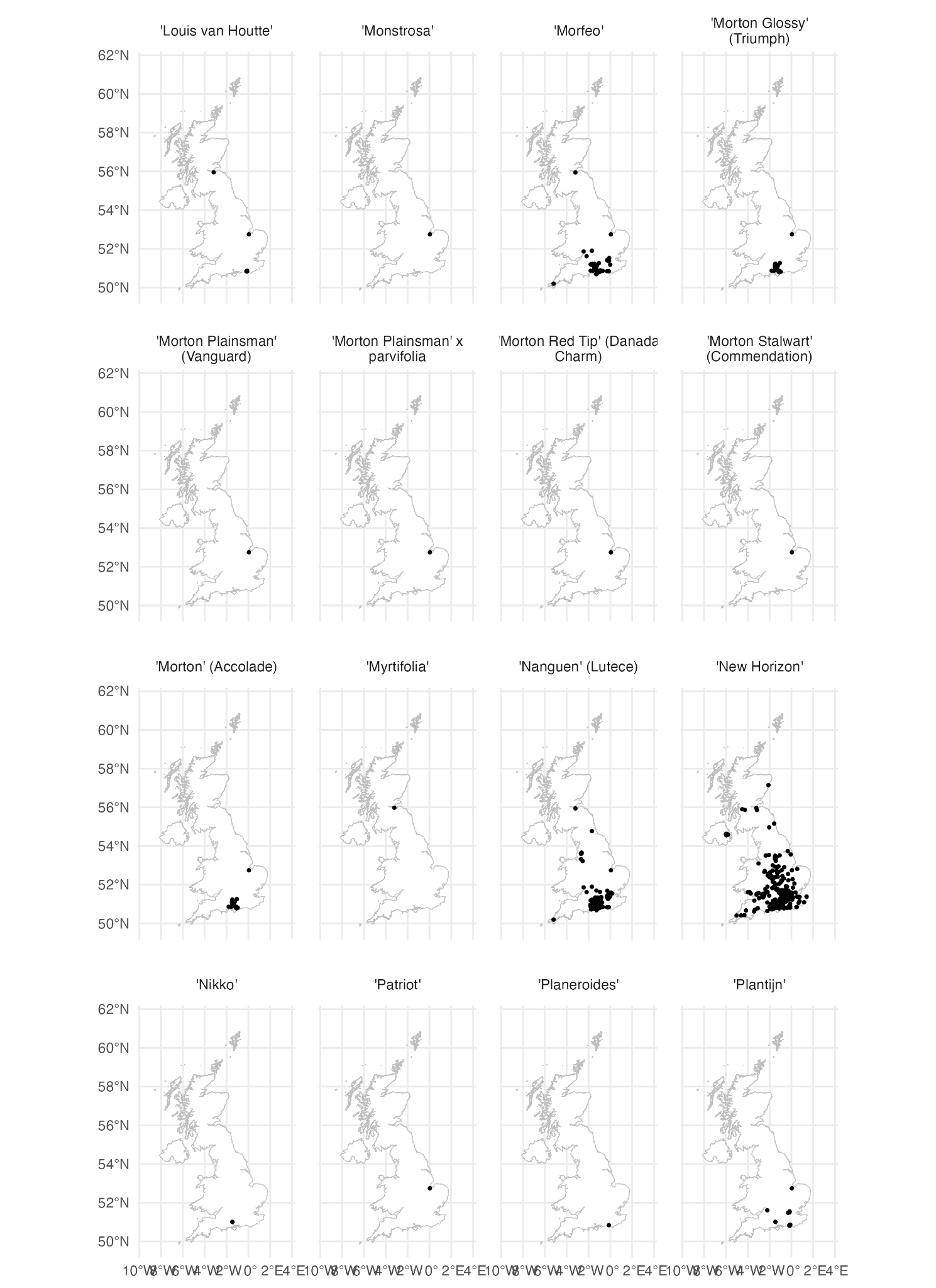

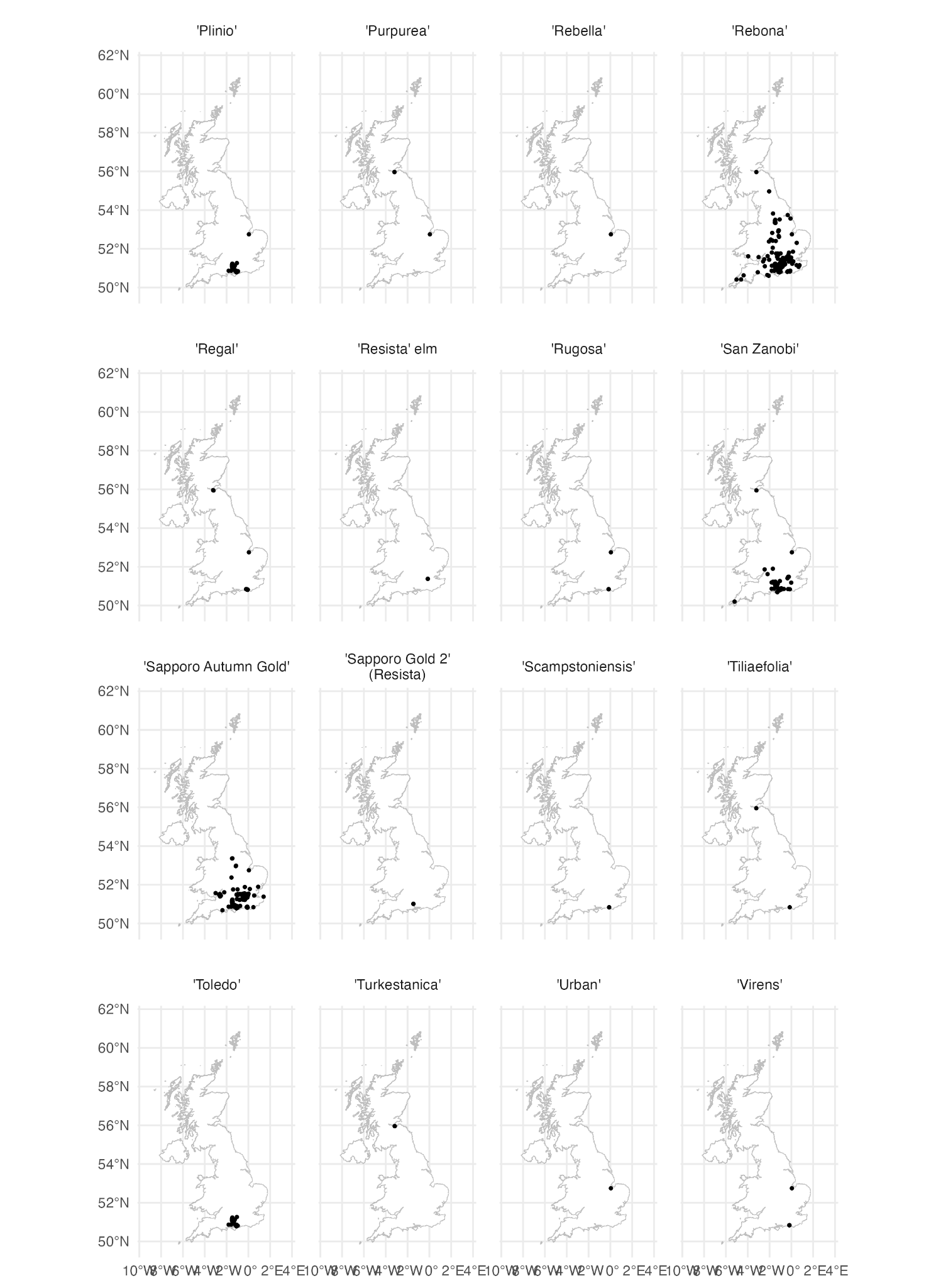

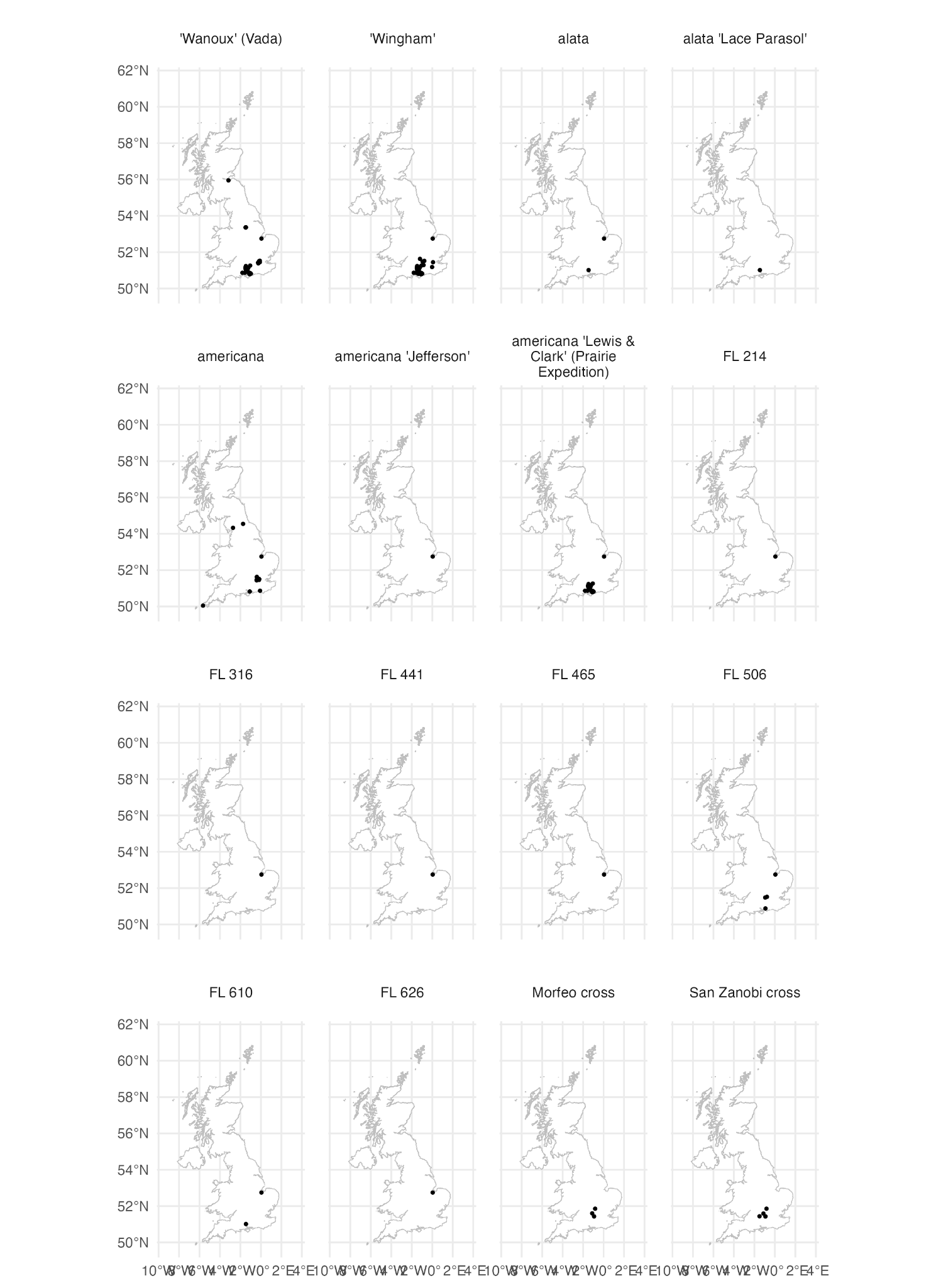

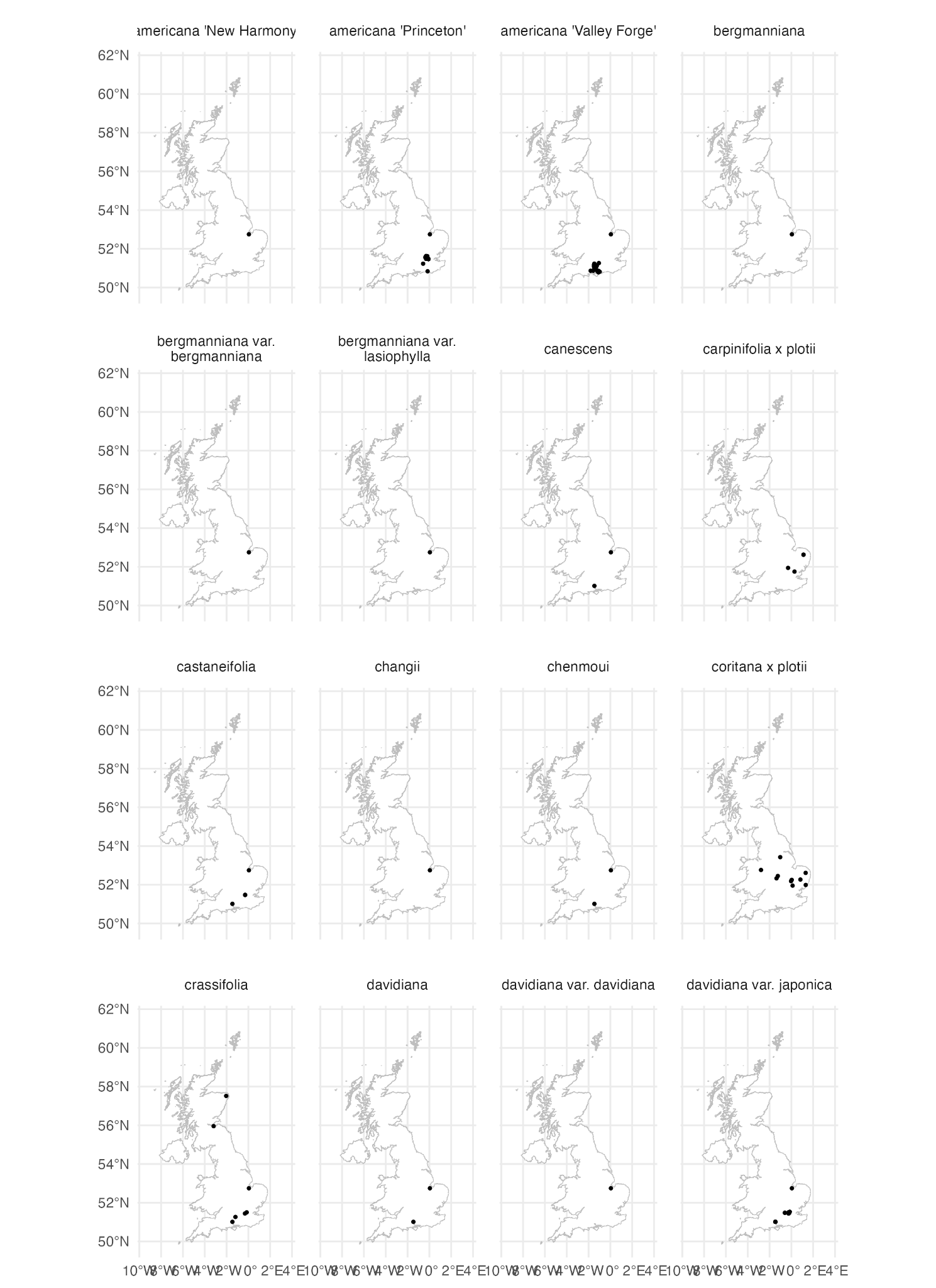

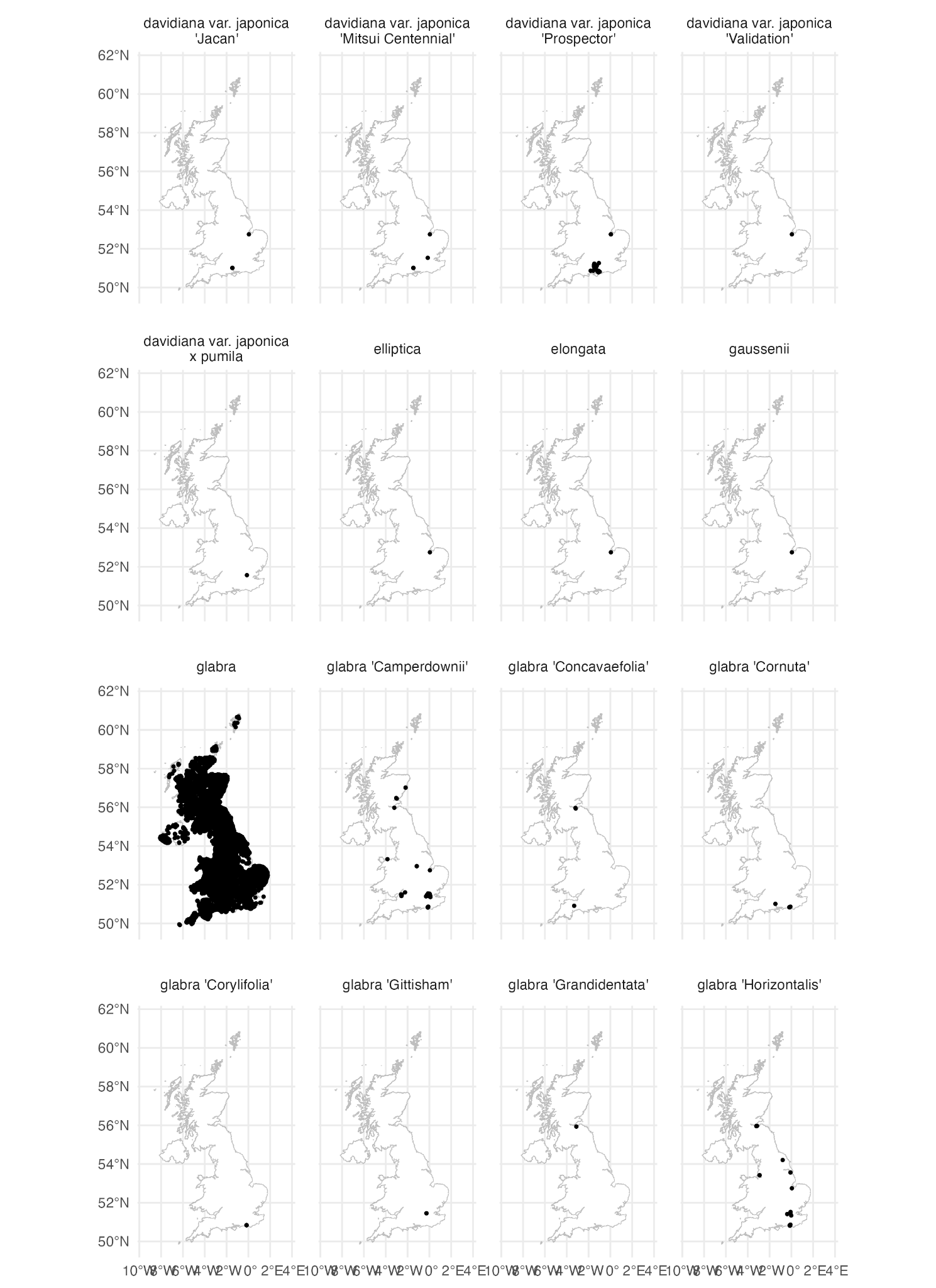

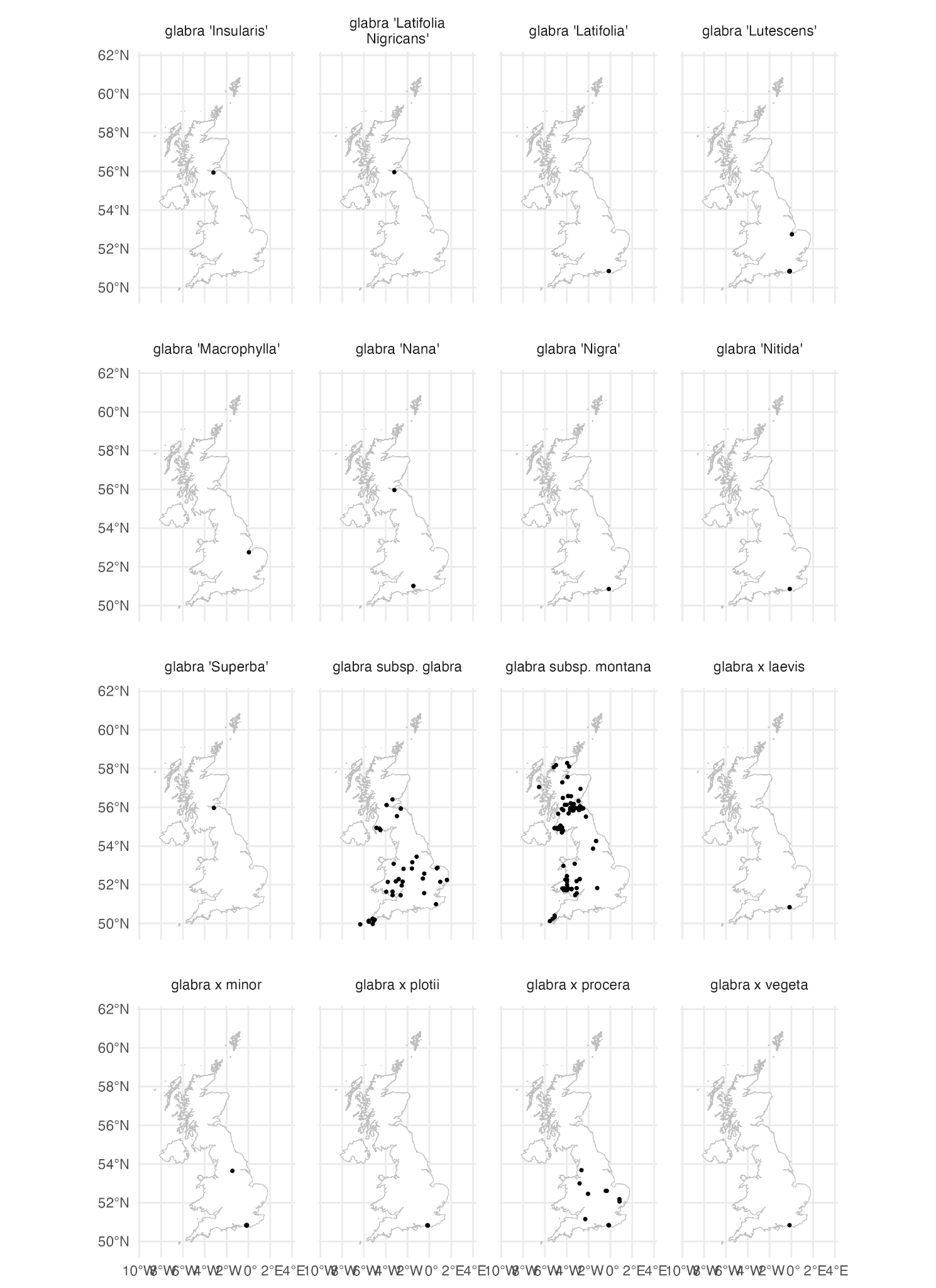

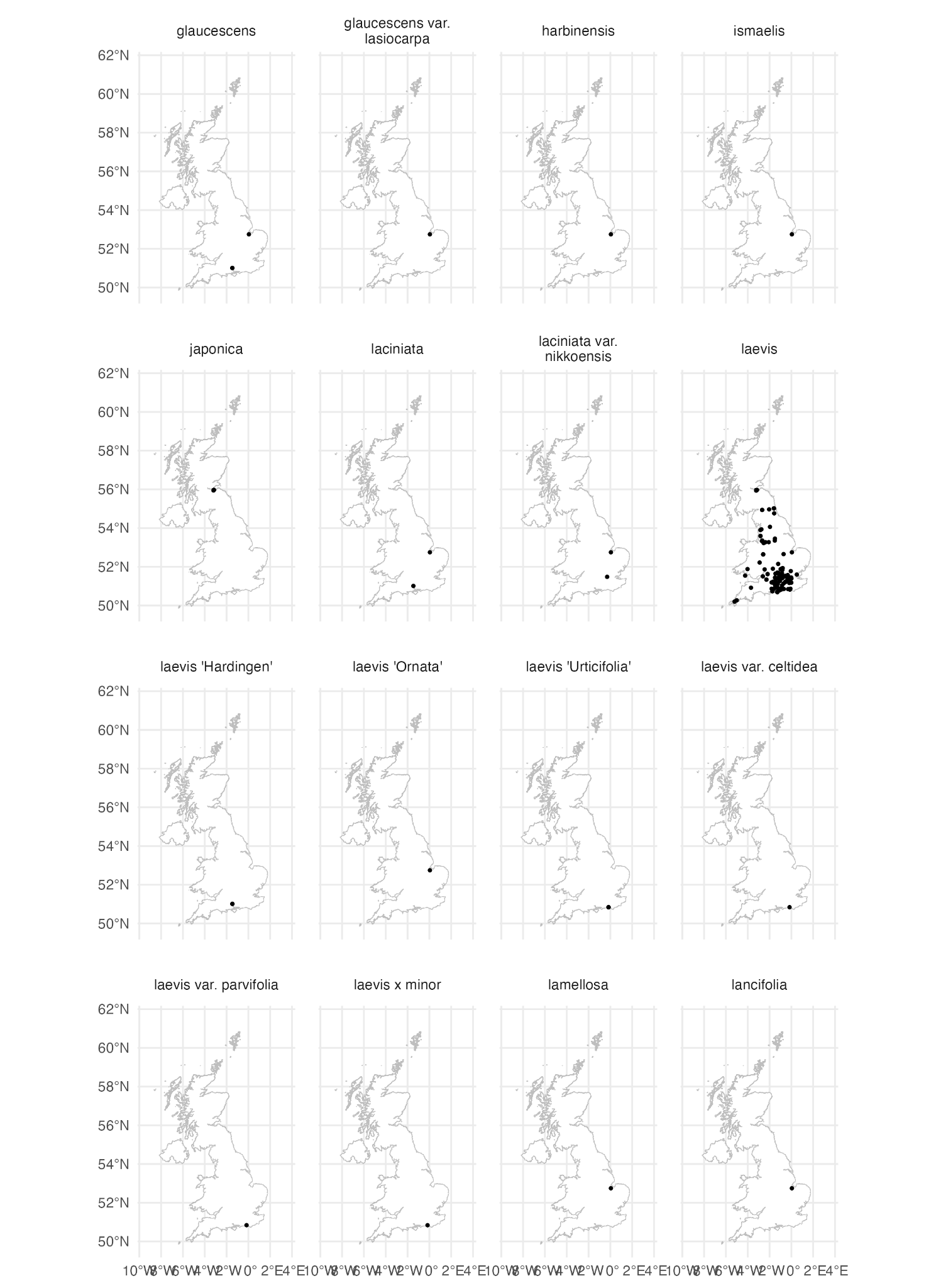

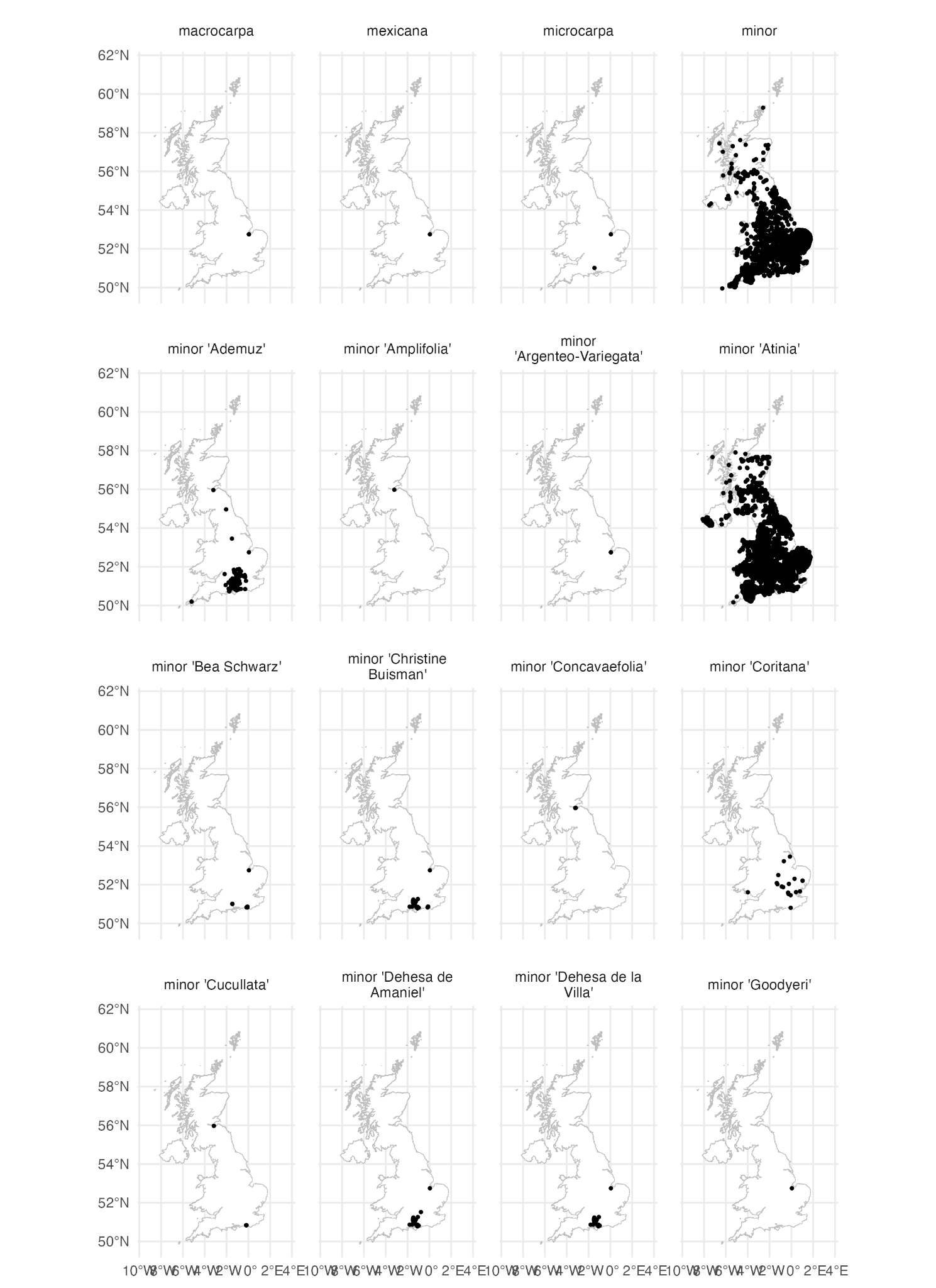

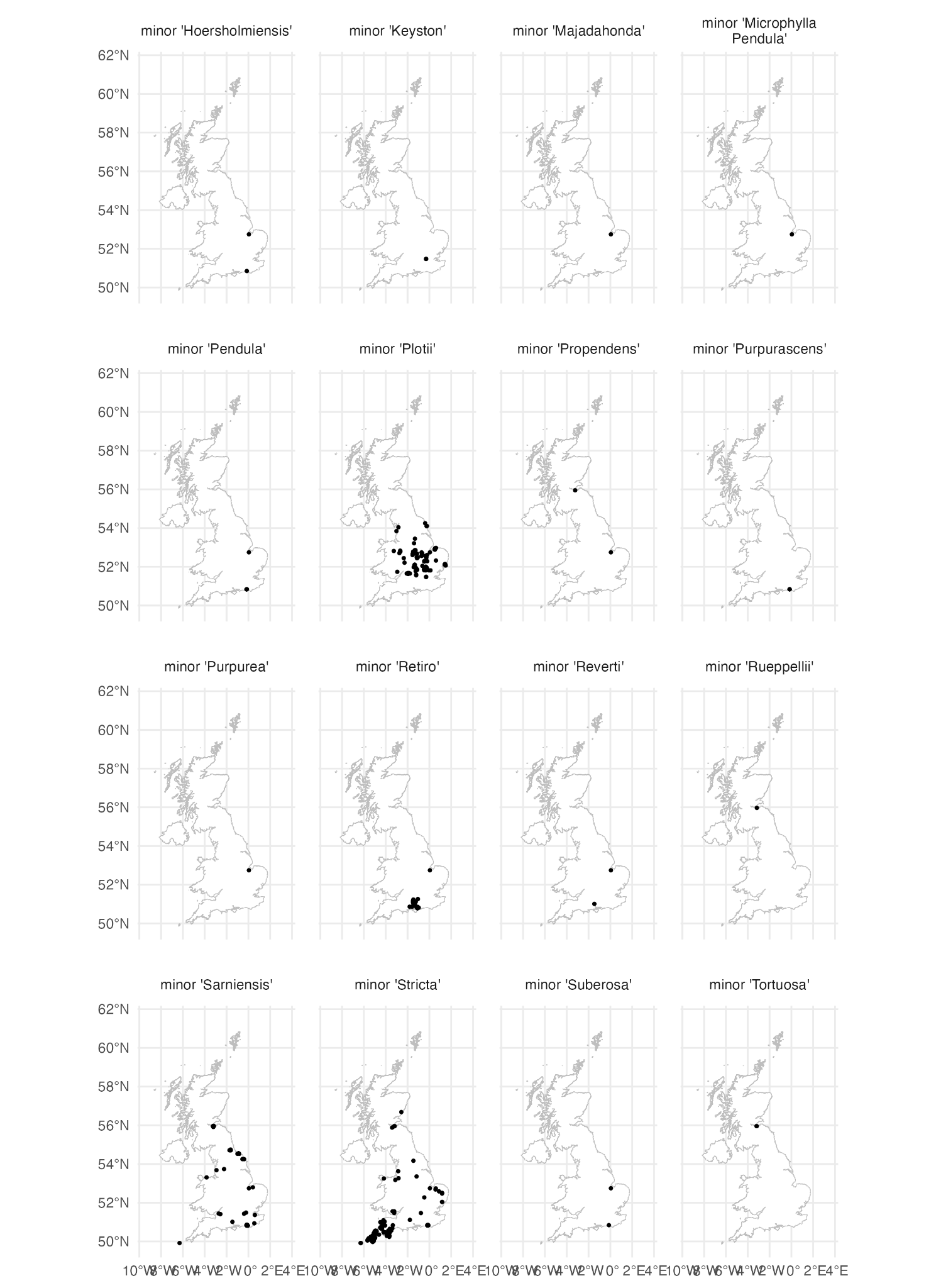

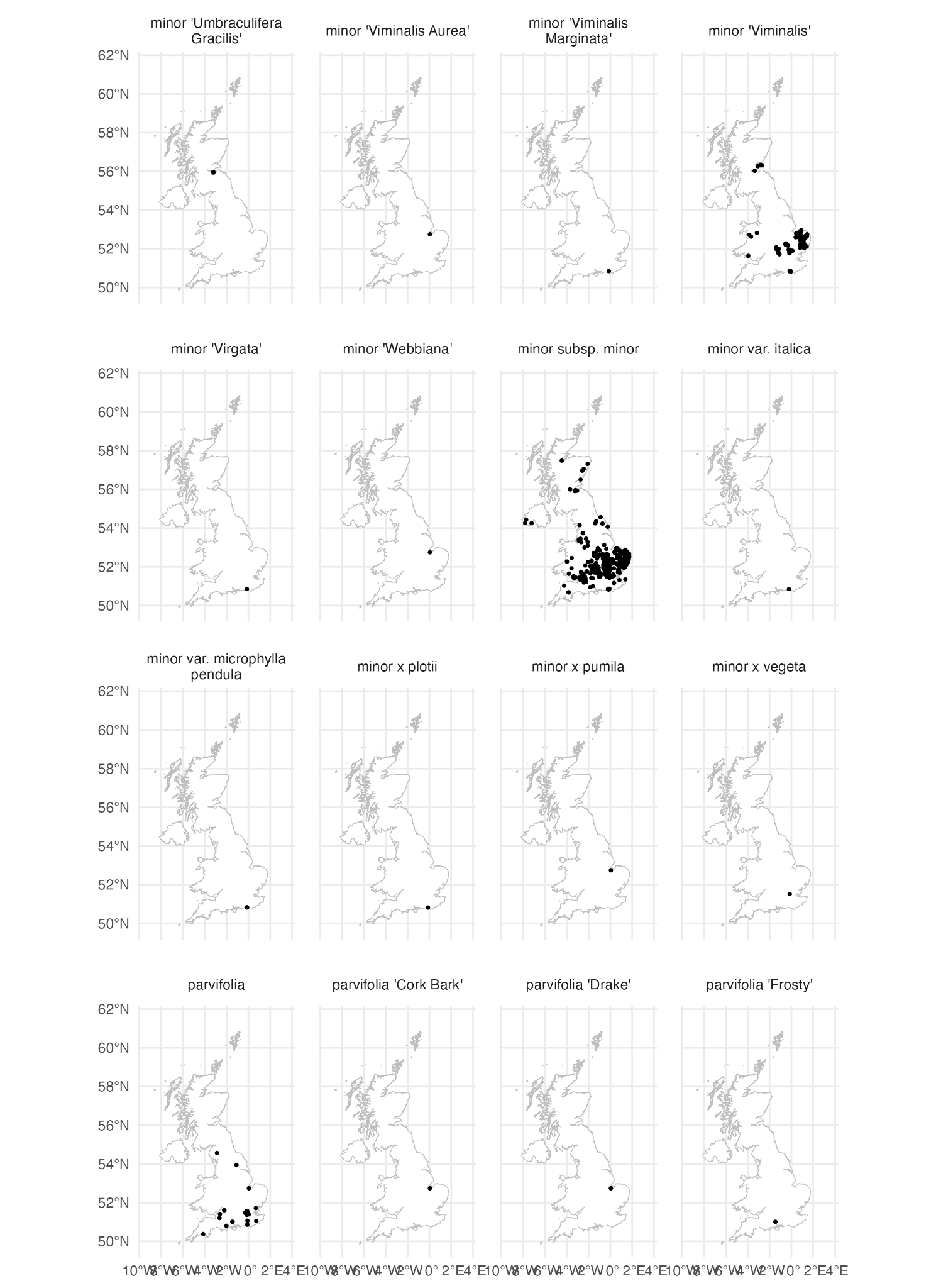

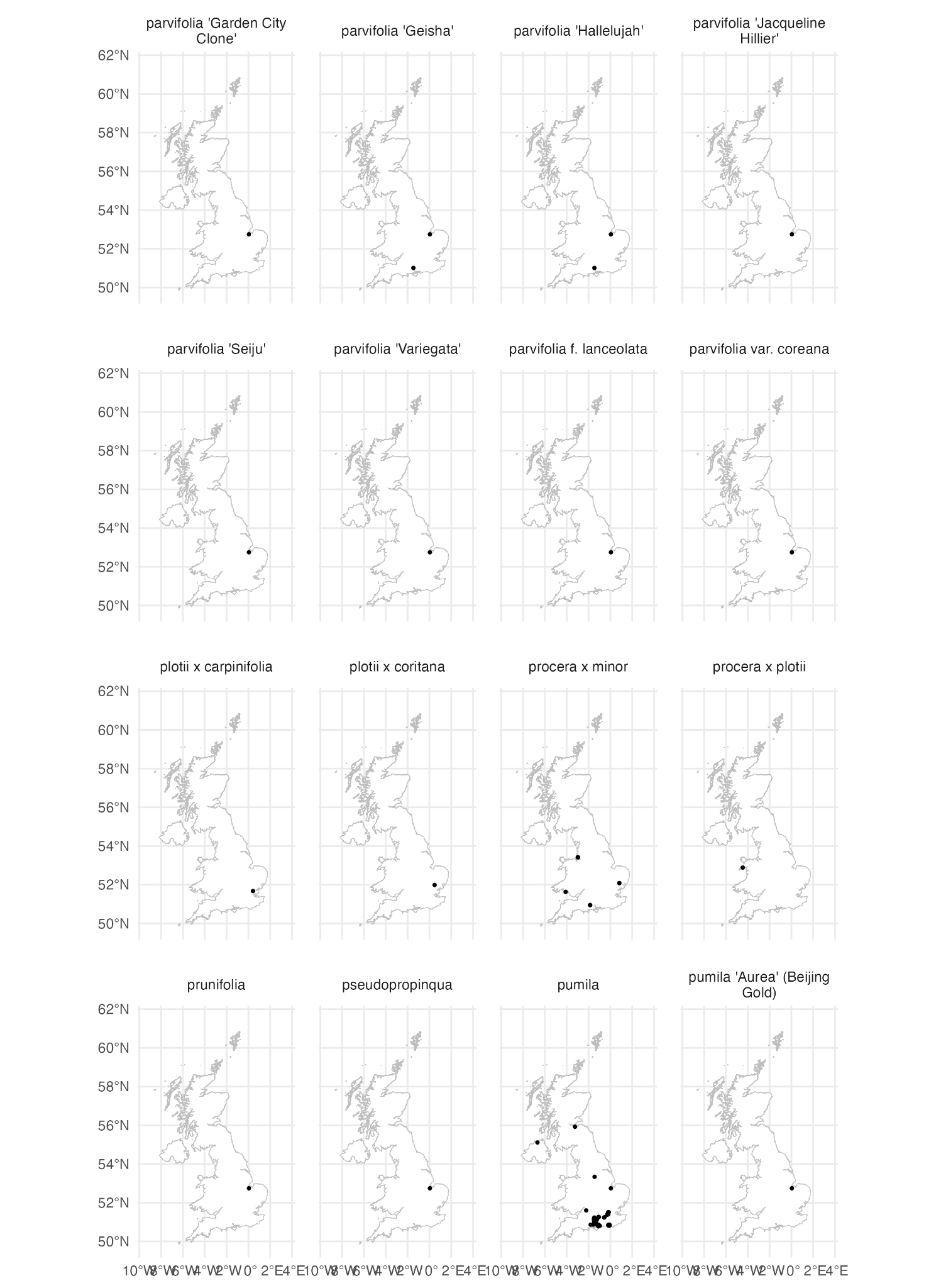

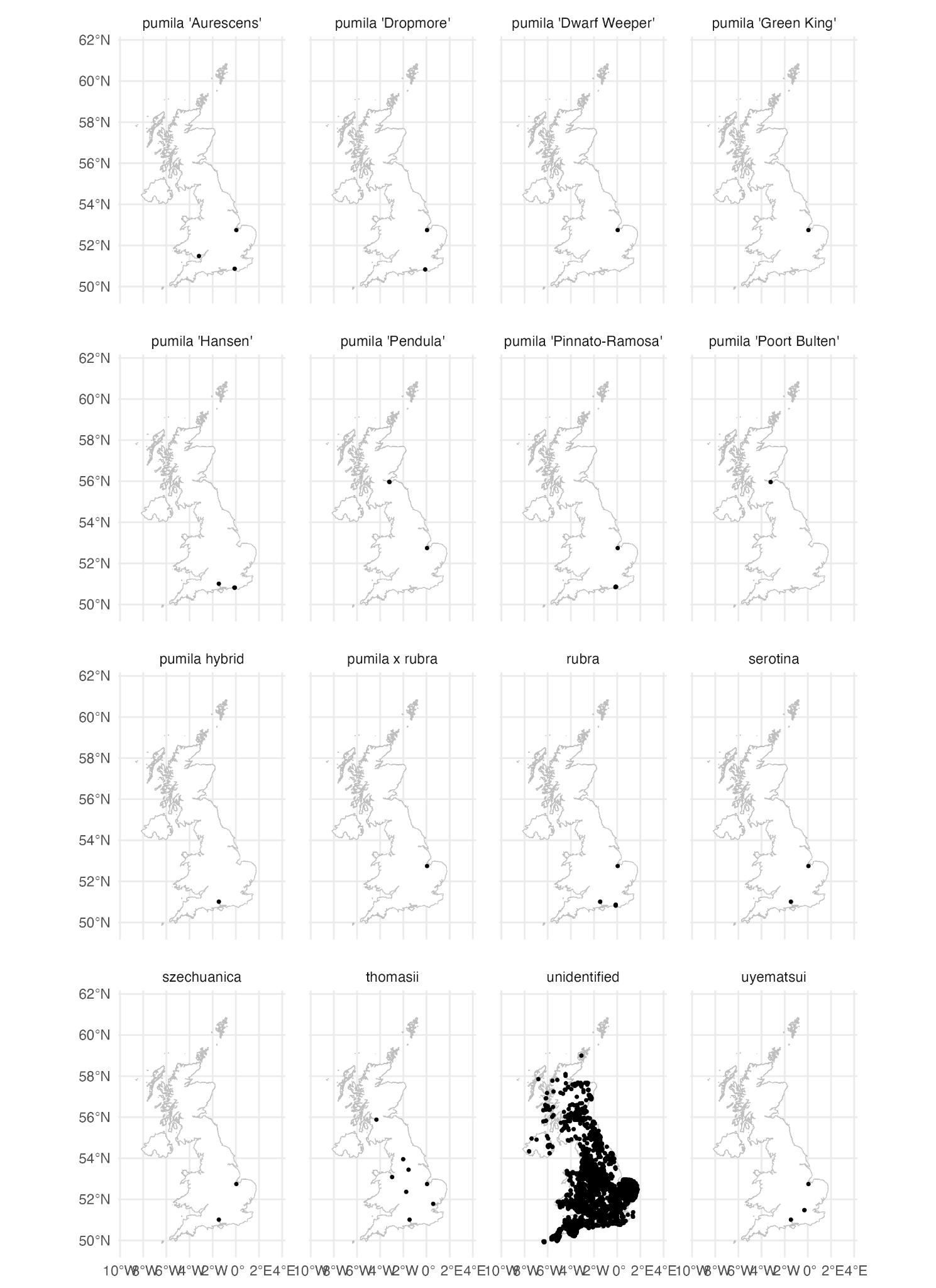

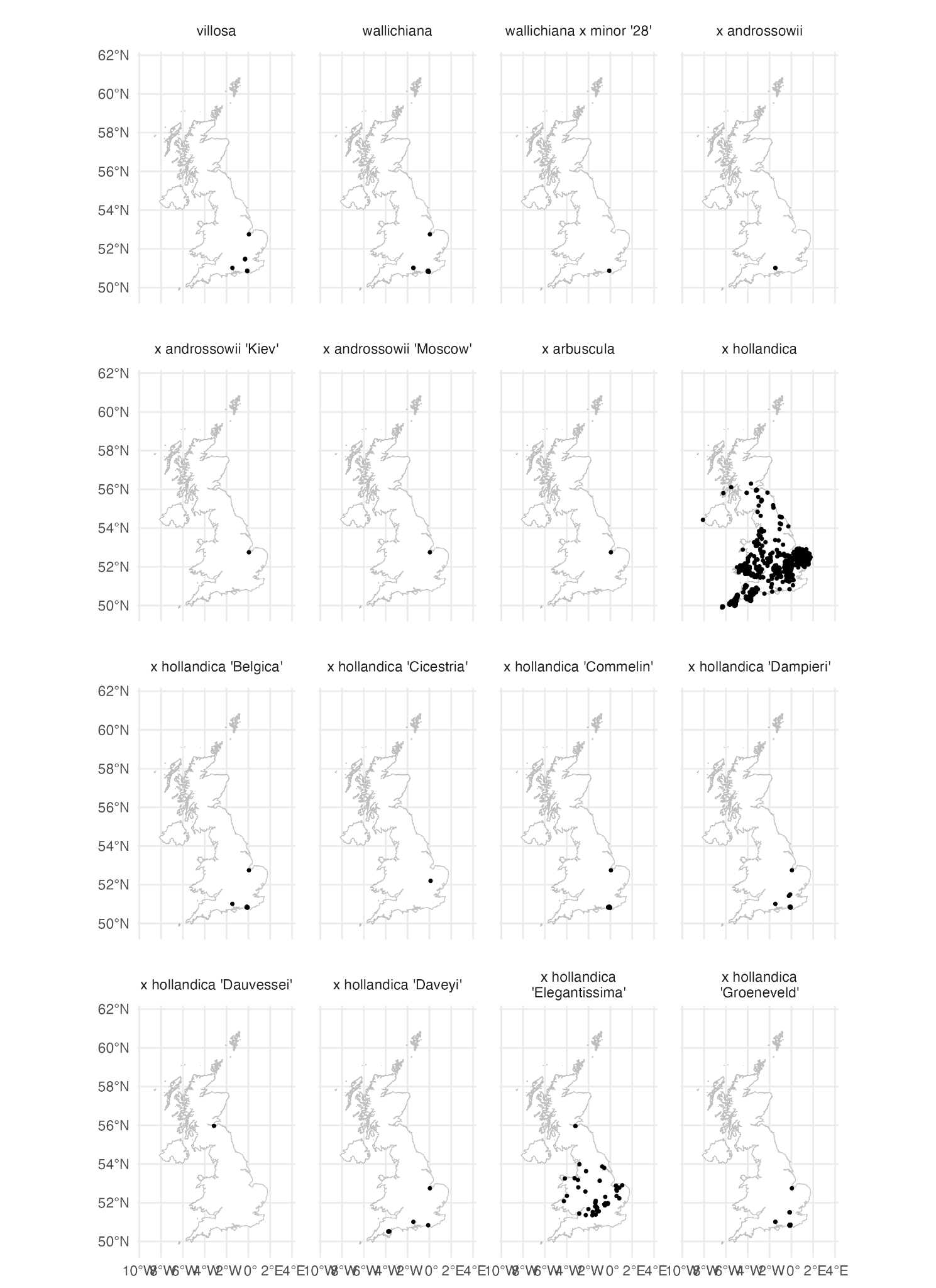

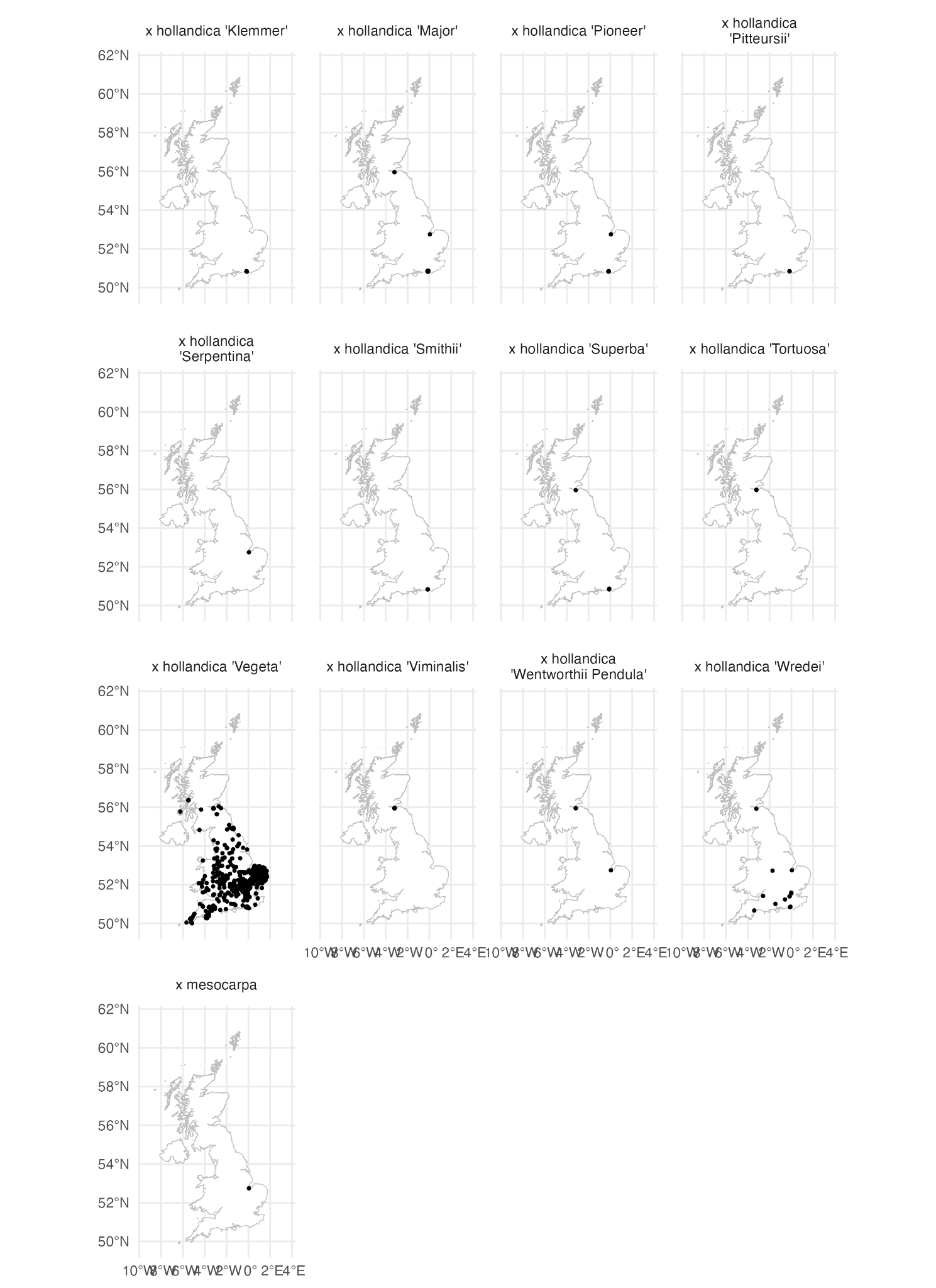

Supplementary Figure: Maps of all cultivars and species that could be georeferenced (and unidentified occurrences)

Supplementary Table: Number of occurrences mapped per elm (identifications are the opinion of the recorders, apart from minimal correction of synonyms).

| **Name** | **Number of occurrences** |
| --- | --- |
| '148' | 26 |
| '202' | 46 |
| '260' | 17 |
| '28' | 9 |
| '297' | 6 |
| '6' | 2 |
| 'Alksuth' | 1 |
| 'Amsterdam' | 2 |
| 'Arno' | 1 |
| 'Atropurpurea' | 1 |
| 'Cathedral' | 1 |
| 'Clusius' | 14 |
| 'Columella' | 89 |
| 'Crispa' | 1 |
| 'Den Haag' | 4 |
| 'Dodoens' | 110 |
| 'Eisele A1' | 23 |
| 'Eisele H1' | 23 |
| 'Europa' | 3 |
| 'Exoniensis' | 25 |
| 'Exoniensis' x wallichiana | 2 |
| 'Fagel' | 2 |
| 'Fastigiata Glabra' | 1 |
| 'Fiorente' | 127 |
| 'Frontier' | 30 |
| 'Fuente Umbria' | 30 |
| 'Homestead' | 8 |
| 'Jacqueline Hillier' | 16 |
| 'Karagatch' | 1 |
| 'Klondike' | 2 |
| 'Lobel' | 430 |
| 'Lombartsii' | 3 |
| 'Louis van Houtte' | 15 |
| 'Monstrosa' | 1 |
| 'Morfeo' | 57 |
| 'Morton Glossy' (Triumph) | 32 |
| 'Morton Plainsman' (Vanguard) | 1 |
| 'Morton Plainsman' x parvifolia | 1 |
| 'Morton Red Tip' (Danada Charm) | 1 |
| 'Morton Stalwart' (Commendation) | 1 |
| 'Morton' (Accolade) | 30 |
| 'Myrtifolia' | 1 |
| 'Nanguen' (Lutece) | 986 |
| 'New Horizon' | 905 |
| 'Nikko' | 2 |
| 'Patriot' | 1 |
| 'Planeroides' | 1 |
| 'Plantijn' | 19 |
| 'Plinio' | 32 |
| 'Purpurea' | 3 |
| 'Rebella' | 1 |
| 'Rebona' | 183 |
| 'Regal' | 19 |
| 'Resista' elm | 2 |
| 'Rugosa' | 3 |
| 'San Zanobi' | 51 |
| 'Sapporo Autumn Gold' | 216 |
| 'Sapporo Gold 2' (Resista) | 2 |
| 'Scampstoniensis' | 2 |
| 'Tiliaefolia' | 2 |
| 'Toledo' | 29 |
| 'Turkestanica' | 2 |
| 'Urban' | 1 |
| 'Virens' | 3 |
| 'Wanoux' (Vada) | 41 |
| 'Wingham' | 54 |
| FL 214 | 1 |
| FL 316 | 1 |
| FL 441 | 1 |
| FL 465 | 1 |
| FL 506 | 4 |
| FL 610 | 2 |
| FL 626 | 1 |
| Morfeo cross | 3 |
| San Zanobi cross | 5 |
| alata | 2 |
| alata 'Lace Parasol' | 1 |
| americana | 17 |
| americana 'Jefferson' | 1 |
| americana 'Lewis & Clark' (Prairie Expedition) | 30 |
| americana 'New Harmony' | 1 |
| americana 'Princeton' | 30 |
| americana 'Valley Forge' | 31 |
| bergmanniana | 1 |
| bergmanniana var. bergmanniana | 1 |
| bergmanniana var. lasiophylla | 1 |
| canescens | 2 |
| carpinifolia x plotii | 4 |
| castaneifolia | 4 |
| changii | 1 |
| chenmoui | 2 |
| coritana x plotii | 10 |
| crassifolia | 7 |
| davidiana | 2 |
| davidiana var. davidiana | 1 |
| davidiana var. japonica | 16 |
| davidiana var. japonica 'Jacan' | 4 |
| davidiana var. japonica 'Mitsui Centennial' | 5 |
| davidiana var. japonica 'Prospector' | 31 |
| davidiana var. japonica 'Validation' | 1 |
| davidiana var. japonica x pumila | 1 |
| elliptica | 1 |
| elongata | 1 |
| gaussenii | 1 |
| glabra | 33520 |
| glabra 'Camperdownii' | 53 |
| glabra 'Concavaefolia' | 3 |
| glabra 'Cornuta' | 7 |
| glabra 'Corylifolia' | 3 |
| glabra 'Gittisham' | 2 |
| glabra 'Grandidentata' | 1 |
| glabra 'Horizontalis' | 20 |
| glabra 'Insularis' | 1 |
| glabra 'Latifolia Nigricans' | 1 |
| glabra 'Latifolia' | 1 |
| glabra 'Lutescens' | 18 |
| glabra 'Macrophylla' | 1 |
| glabra 'Nana' | 2 |
| glabra 'Nigra' | 1 |
| glabra 'Nitida' | 1 |
| glabra 'Superba' | 1 |
| glabra subsp. glabra | 44 |
| glabra subsp. montana | 120 |
| glabra x laevis | 2 |
| glabra x minor | 17 |
| glabra x plotii | 3 |
| glabra x procera | 14 |
| glabra x vegeta | 1 |
| glaucescens | 3 |
| glaucescens var. lasiocarpa | 1 |
| harbinensis | 1 |
| ismaelis | 1 |
| japonica | 4 |
| laciniata | 2 |
| laciniata var. nikkoensis | 2 |
| laevis | 303 |
| laevis 'Hardingen' | 2 |
| laevis 'Ornata' | 1 |
| laevis 'Urticifolia' | 3 |
| laevis var. celtidea | 1 |
| laevis var. parvifolia | 1 |
| laevis x minor | 1 |
| lamellosa | 1 |
| lancifolia | 1 |
| macrocarpa | 2 |
| mexicana | 1 |
| microcarpa | 2 |
| minor | 6462 |
| minor 'Ademuz' | 640 |
| minor 'Amplifolia' | 1 |
| minor 'Argenteo-Variegata' | 2 |
| minor 'Atinia' | 22334 |
| minor 'Bea Schwarz' | 24 |
| minor 'Christine Buisman' | 37 |
| minor 'Concavaefolia' | 2 |
| minor 'Coritana' | 19 |
| minor 'Cucullata' | 3 |
| minor 'Dehesa de Amaniel' | 31 |
| minor 'Dehesa de la Villa' | 31 |
| minor 'Goodyeri' | 1 |
| minor 'Hoersholmiensis' | 2 |
| minor 'Keyston' | 3 |
| minor 'Majadahonda' | 1 |
| minor 'Microphylla Pendula' | 1 |
| minor 'Pendula' | 4 |
| minor 'Plotii' | 115 |
| minor 'Propendens' | 2 |
| minor 'Purpurascens' | 3 |
| minor 'Purpurea' | 1 |
| minor 'Retiro' | 30 |
| minor 'Reverti' | 2 |
| minor 'Rueppellii' | 1 |
| minor 'Sarniensis' | 167 |
| minor 'Stricta' | 285 |
| minor 'Suberosa' | 2 |
| minor 'Tortuosa' | 2 |
| minor 'Umbraculifera Gracilis' | 3 |
| minor 'Viminalis Aurea' | 1 |
| minor 'Viminalis Marginata' | 1 |
| minor 'Viminalis' | 84 |
| minor 'Virgata' | 1 |
| minor 'Webbiana' | 1 |
| minor subsp. minor | 632 |
| minor var. italica | 1 |
| minor var. microphylla pendula | 2 |
| minor x plotii | 1 |
| minor x pumila | 1 |
| minor x vegeta | 1 |
| parvifolia | 66 |
| parvifolia 'Cork Bark' | 1 |
| parvifolia 'Drake' | 1 |
| parvifolia 'Frosty' | 1 |
| parvifolia 'Garden City Clone' | 1 |
| parvifolia 'Geisha' | 2 |
| parvifolia 'Hallelujah' | 2 |
| parvifolia 'Jacqueline Hillier' | 1 |
| parvifolia 'Seiju' | 1 |
| parvifolia 'Variegata' | 1 |
| parvifolia f. lanceolata | 1 |
| parvifolia var. coreana | 1 |
| plotii x carpinifolia | 1 |
| plotii x coritana | 1 |
| procera x minor | 7 |
| procera x plotii | 1 |
| prunifolia | 1 |
| pseudopropinqua | 1 |
| pumila | 99 |
| pumila 'Aurea' (Beijing Gold) | 1 |
| pumila 'Aurescens' | 3 |
| pumila 'Dropmore' | 2 |
| pumila 'Dwarf Weeper' | 1 |
| pumila 'Green King' | 1 |
| pumila 'Hansen' | 6 |
| pumila 'Pendula' | 4 |
| pumila 'Pinnato-Ramosa' | 7 |
| pumila 'Poort Bulten' | 1 |
| pumila hybrid | 1 |
| pumila x rubra | 1 |
| rubra | 6 |
| serotina | 2 |
| szechuanica | 2 |
| thomasii | 8 |
| unidentified | 10421 |
| uyematsui | 4 |
| villosa | 8 |
| wallichiana | 61 |
| wallichiana x minor '28' | 1 |
| x androssowii | 2 |
| x androssowii 'Kiev' | 1 |
| x androssowii 'Moscow' | 1 |
| x arbuscula | 1 |
| x hollandica | 1020 |
| x hollandica 'Belgica' | 38 |
| x hollandica 'Cicestria' | 1 |
| x hollandica 'Commelin' | 61 |
| x hollandica 'Dampieri' | 16 |
| x hollandica 'Dauvessei' | 1 |
| x hollandica 'Daveyi' | 7 |
| x hollandica 'Elegantissima' | 51 |
| x hollandica 'Groeneveld' | 52 |
| x hollandica 'Klemmer' | 5 |
| x hollandica 'Major' | 66 |
| x hollandica 'Pioneer' | 9 |
| x hollandica 'Pitteursii' | 3 |
| x hollandica 'Serpentina' | 1 |
| x hollandica 'Smithii' | 2 |
| x hollandica 'Superba' | 4 |
| x hollandica 'Tortuosa' | 2 |
| x hollandica 'Vegeta' | 1054 |
| x hollandica 'Viminalis' | 3 |
| x hollandica 'Wentworthii Pendula' | 3 |
| x hollandica 'Wredei' | 16 |
| x mesocarpa | 1 |
